# PI 3-kinase isoform p110α mediated FOXO1 inactivation protects against aortic aneurysm formation

**DOI:** 10.1101/2022.12.01.518561

**Authors:** Marius Vantler, Maximilian Schorscher, Eva M. Berghausen, Mario Zierden, Jasper Spitzer, Joseph B. Moore, Dickson W.L. Wong, Zhaolong Li, Max Wißmüller, Leoni Gnatzy-Feik, David Unnersjö-Jess, Laura-Marie A. Zimmermann, Maria Grandoch, Dennis Mehrkens, Matti Adam, Xinlei Zhao, Susanne V. Schmidt, Maximilian Billmann, Margarete Odenthal, Gerhard Sengle, Chanil Valasarajan, Soni Pullamsetti, Peter Boor, Lars Maegdefessel, Stephan Baldus, Stephan Rosenkranz

## Abstract

**Aims:** Abdominal aortic aneurysms (AAAs) are characterized by loss of contractile smooth muscle cells (SMCs). Regenerative approaches for AAAs should therefore aim at enriching aortic SMC content. We explored the role of PI 3-kinase isoform p110α, a crucial regulator of cellular proliferation, survival and phenotype, for vascular integrity and AAA disease.

**Methods and results:** Mice lacking p110α or FOXO1 in SMCs (sm-p110α^−/−^, sm-FOXO1^−/−^) were exposed to the porcine pancreatic elastase (PPE) AAA model. Phenotypic alterations were examined by ultrasound, (immuno)histochemistry, transmission electron microscopy, functional and cellular assays, and single cell RNA-sequencing (scRNA-seq) in tamoxifen-inducible ism-p110α^−/−^-eYFP lineage tracing mice. PPE treatment of sm-p110α^−/−^ mice yielded aggravated AAA formation compared to wild-type controls. AAA onset in sm-p110α^−/−^ aortae was facilitated by a subtle preexisting basal phenotype displaying smaller media area, impaired aortic wall structure, and diminished responsiveness to vasodilators. During AAA progression, PPE treated sm-p110α^−/−^ aortae were characterized by impaired vascular remodeling, reduced SMC content and medial cell proliferation, and increased elastic fiber breaks and ECM degradation. scRNA-seq revealed decreased content of proliferative and reparative SMC clusters and increased inflamed SMCs in ism-p110α^−/−^-eYFP mice. p110α deficient SMCs displayed impaired SMC proliferation, and production of elastin fibers. This phenotype was rescued by FOXO1 inactivation in p110α^−/−^ SMCs. FOXO1 deficiency in SMCs protected against AAA in PPE-treated sm-FOXO1^−/−^ mice. Pharmacological activation of p110α ameliorated AAA in wild type mice but not in sm-FOXO1^−/−^ mice. Relevance of p110α/FOXO1 signaling in human AAA disease was indicated in patient samples.

**Conclusions:** p110α-mediated inactivation of FOXO1 in SMCs protects against AAA disease and contributes to maintaining aortic wall homoeostasis. p110α signaling promotes transition into proliferating and reparative SMC phenotypes while attenuating de-differentiation into inflammatory SMC subtypes. Our findings identify (pharmacological) activation of p110α signaling in SMCs as a new therapeutic strategy to promote prevention and regeneration of AAA disease.

**Translational Perspective:** AAAs account for ∼80% of all aneurysms, and rupture is associated with significant mortality (∼85%). No effective pharmacological approach has been identified that can limit the progression or rupture of AAAs. A major cause of AAA formation is the loss of contractile SMCs. We identified the p110α/AKT/FOXO1 pathway as an attractive therapeutic target to increase SMC content without inducing detrimental SMC de-differentiation in AAAs. Activation of p110α signaling in SMCs may protect patients from AAA progression and initiate regeneration of the diseased aortic segment. Our findings may also raise potential safety concerns for current approaches aimed at p110α inhibition for cancer therapy.

**Graphical Abstract:** Analysis of (pharmacological) activation of p110α signaling in sm-p110α^+/+^ and sm-p110α^−/−^ mice revealed p110α dependent FOXO1 inactivation in SMCs which maintains SMC functionality (proliferation, ECM production) contributing to aortic wall structure and function as well as protection against AAA progression in PPE treated mice. GF: growth factor; RTK: receptor tyrosine kinase; SMC: aortic smooth muscle cell; ECM: extracellular matrix; AAA: abdominal aortic aneurysm.

## Introduction

Abdominal aortic aneurysms (AAAs), accounting for approximately 80% of all aneurysms, are pathological dilations of the abdominal aorta, which increases in diameter to ≥30 mm (or 1.5 times its normal size).^1,2^ AAAs are most commonly found in the infrarenal abdominal aorta, with a global prevalence of 0.92% among people aged 30-79.^3,4^ Unless they rapidly increase in size, rupture, or thromboembolize into the distal arteries, AAAs remain clinically asymptomatic.^5^ Fatal outcomes due to rupture occur when the intraluminal pressure and wall stress exceed the strength of the aortic wall. The mortality rate is high (∼85%),^13^ and it is estimated that ruptured AAAs are responsible for nearly 15,000 deaths in the United States per year.^6^ No effective approach has been identified to date that can limit AAA progression or the risk of rupture.^7^ Hence, the only available treatment option remains surgical intervention, which includes open or endovascular aortic repair.^2^ To identify novel therapeutic targets, it is essential to develop a better understanding of the cellular mechanisms and regulatory networks driving AAA development and progression.

AAAs feature alterations of the aortic wall structure, including thinning of the media due to loss of vascular smooth muscle cells (SMCs) and degradation of the extracellular matrix (ECM).^5^ SMCs are the predominant cell type within the aortic wall and have a remarkable phenotypic plasticity. Important functions in both healthy and disease conditions include apoptosis, proliferation, phenotypic switching, contractility, as well as ECM production and degradation. The aortic ECM architecture is assembled by supra-molecular networks composed of elastin and collagen, which play essential roles in maintaining vessel homeostasis. Damage or inappropriate remodeling of the ECM contributes to the initiation and progression of AAA disease.^8^ Stress signals within the aortic wall, including oxidative stress, inflammatory cell infiltration, dysregulated metabolic homeostasis, and (epi)genetic changes, promote SMC death.^9^ To compensate for SMC loss, SMCs switch from contractile to de-differentiated proliferative phenotypes^10,11^ that resemble inflammatory cells,^12^ fibroblasts,^13^ and osteogenic cells.^14^ Recent evidence suggests that SMC phenotypic alterations are a common feature of AAA disease,^15–17^ and that diverse phenotypes may have distinct effects on AAA progression. Whereas proliferative phenotypes may serve aortic repair and compensate for SMC loss during AAA development, the switch to inflammatory phenotypes may promote aortic dysfunction and degeneration. The molecular mechanisms underlying these changes have yet to be identified. However, understanding these mechanisms is necessary for developing strategies to maintain or restore aortic wall homeostasis.

Our previous work revealed that the catalytic class IA phosphatidylinositol 3′-kinase (PI3K) isoform p110α is a crucial driver of SMC proliferation and migration, and contributes to vascular diseases including neointima formation after balloon injury and pulmonary vasculopathy.^18,19^ Class IA PI3Ks signal downstream of receptor tyrosine kinases (RTKs), and consist of a regulatory p85 subunit and one of three distinct 110-kDa catalytic subunits (p110α, p110β, p110δ). Class IA PI3Ks are lipid kinases generating phosphatidyinositol-3,4,5-trisphosphate (PIP_3_),^20–22^ which serve as a docking site for downstream signaling molecules such as AKT, which is considered the major transducer of PI3K signal relay. AKT phosphorylates numerous target proteins including major regulators of metabolism, mitogenesis, and apoptosis including Forkhead Box O (FOXO) transcription factors.^23^ Recently, the development of a pharmacological activator of p110α was reported, which provides cardioprotection from ischemia-reperfusion injury.^24^

Although p110α signaling is implicated in various vascular diseases, its potential physiological functions in maintenance of vascular integrity as well as its pathological role in AAA disease are largely unknown. Since p110α represents a critical mediator of cell proliferation, it is assumed to play a significant role during AAA development.^9^ Herein, we aimed to elucidate the role of p110α in SMC proliferation, phenotypic modulation, and ECM homeostasis in the aortic wall and its impact on AAA disease.

## Materials and Methods

Comprehensive methodological details are presented in the Supplemental Material.

### Generation of smooth muscle–specific p110α- and FOXO1-deficient mice

Smooth muscle-specific p110α– and FOXO1-deficient mice were generated by cross-breeding homozygous *Pik3ca^fl/fl^* or *Foxo1^fl/fl^*mice^25,26^ with *Sm22-Cre* mice (B6.Cg-Tg[Tagln-Cre]1Her/J, The Jackson Laboratory), which express Cre recombinase under the control of the mouse transgelin (smooth muscle protein 22-alpha) promoter. *SM22-Cre^+/−^/Pik3ca^fl/+^*or *Sm22-Cre^+/−^/FOXO1^fl/+^* mice were crossed to homozygous *Pik3ca^fl/fl^* or *Foxo1^fl/fl^* mice, respectively, to generate *Sm22-Cre^−^/Pik3ca^fl/fl^* (sm-p110α^+/+^) and *Sm22-Cre^+/−^/Pik3ca^fl/fl^* (sm-p110α^−/−^)^19^ as well as *Sm22-Cre^−^/Foxo1^fl/fl^* (sm-FOXO1^+/+^) and *Sm22-Cre^+^/Foxo1^fl/fl^* (sm-FOXO^−/−^) littermates. Genotyping was performed by PCR using ear punches.

Male sex is one of the most important risk factors of AAA disease (∼4–6 times more prevalent in men compared with women). Therefore, we focused in this study on male mice.

### PPE perfusion model

To induce murine AAAs, the PPE infusion model was performed as previously described.^27^ Briefly, adult male mice (weight approx. 25-30 g) first received analgesia 30 minutes before surgery via subcutaneous injection of buprenorphine (0.1 mg/kg body weight, 1 ml Temgesic® (buprenorphine 0.3 mg/ml)) mixed with 5 ml 0.9% NaCl solution. Anesthesia is then induced by inhalation with 3.5% isoflurane and subsequently maintained continuously with 1-2% as needed. The animals are secured on a heating pad to protect them from cooling down, with regular manual checks to prevent overheating. The animals receive Eye Care Gel (Remend Cornea®) to protect them from drying out. The anesthetized mice undergo a median laparotomy and the abdominal aorta is isolated at the level of the bifurcation and the left renal vein. All aortic branches within 1.0 cm of the bifurcation are ligated. Temporary ligatures are then placed around the proximal and distal aorta. An aortomy in the region of the bifurcation is then performed. A syringe pump was used to infuse the aorta 100mmHg with saline or saline solution containing 0.04 U / ml of type I porcine pancreatic elastin (Sigma-Aldrich). The perfusion catheter is then removed and the aortomy was repaired without narrowing of the aorta. Analgesia with buprenorphine (0.1 mg/kg body weight) is administered twice daily for up to 3 days post-op. In addition, the mice are supplied with buprenorphine continuously via their drinking water (0,009mg/ml). No PPE treated animal died from aortic rupture. However, approximately 10% of PPE exposed mice died during surgery. All included mice were sacrificed after the indicated period of time following PPE surgery via cervical dislocation. At the end of the experiment, mice were sacrificed. The induced AAA aortic segment (area between the left renal artery and the bifurcation) were harvested at 0, 3, 7, and 28 days post-surgery. Samples were snap-frozen in liquid nitrogen and then stored at −80°C pending further processing.

### Aortic diameter measurements by ultrasound imaging

At baseline, and 7, 14, 21, and 28 days after aneurysm induction, ultrasound imaging was performed to assess the abdominal aortic diameter. Mice were anesthetized with isoflurane 3-4% for induction and then with continuously delivered 2% isoflurane gas inhalation. Ultrasound gel was applied to the depilated skin of the abdomen and imaging was performed with a Vevo3100 imaging system (VisualSonics). B-mode, M-mode and EKV recordings were performed using a MX550D linear array transducer (25-55 MHz, Centre Transmit: 40 MHz, Axial Resolution: 40 μm) with a frame rate of 230–400 frames/s. All ultrasound images were analyzed using the Vevo 3100-software and parameters like aortic diameter were calculated.

### Statistics

All data are expressed as means ± SD. Statistical analysis was performed with the Prism software package (GraphPad Version 10). Differences between two means were assessed by an unpaired t-test. Unless otherwise noted in the figure legends, differences among multiple means were assessed by 2-way ANOVA followed by Bonferroni correction, or uncorrected Fisher’s LSD as well as repeated measures 2-way ANOVA followed by Bonferroni correction (≥2 × 3 factorial design), or uncorrected Fisher’s LSD (2 × 2 factorial design). We used mixed-effects model in case of missing values. Fisher’s exact test was used to determine the significance of associations between categorical variables. P-values <0.05 were considered significant.

### Study approval

The study adhered to the European Parliament Directive 2010/63/EU on the protection of animals used for scientific purposes. Animal experiments were performed in adherence with the National Institutes of Health Guidelines on the Use of Laboratory Animals and were approved by the local animal care committee (Landesamt für Verbraucherschutz und Ernährung (department 2.2) Nordrhein-Westfalen, Recklinghausen, Germany) and the District Council Cologne (approval numbers 81-02.04.2020.A249, 81-02.04.2023.A029, and §4.24.012). For human tissue samples, the study protocol for tissue donation was approved by the Ethics Committee of the University of Munich in accordance with national law and with Good Clinical Practice/International Conference on Harmonization guidelines. Written informed consent was obtained from each patient or the patient’s next of kin.

## Results

### p110α deficiency in SMCs promotes AAA formation in PPE treated mice

To investigate the impact of p110α in SMCs on AAA formation, we exposed sm-p110α^−/−^ mice to the PPE model of AAA formation (**Figure 1A**). Analysis of sm-p110α^−/−^ mice and sm-p110α^+/+^ controls revealed significantly increased AAA growth in sm-p110α^−/−^ mice compared to wild-type controls 28 days after PPE surgery (**Figure 1B,C**), displaying a higher incidence of AAA disease (67% versus 30%), and a more pronounced increase of the aortic diameter (**Figure 1D,E, Supplementary Figure 1**). AAA induction in the infrarenal segment of the aorta led to activation of PI3K signaling in the medial compartment of wild-type mice as monitored by phosphorylation of AKT (**Figure 1F**). This response was diminished in sm-p110α^−/−^ mice, demonstrating the importance of p110α for AAA associated PI3K signaling in medial SMCs. Furthermore, PPE induced AAA formation was accompanied by infiltration of MOMA2^+^ macrophages into the aortic media, specifically in sm-p110α^−/−^ mice (**Figure 1G**). These data indicate that p110α deficiency in SMCs promotes AAA formation and vascular inflammation. We hypothesized that lack of p110α expression could affect AAA formation in two ways: p110α deficiency might promote onset of AAA disease by predisposing sm-p110α^−/−^ mice to structural alterations in the aortic wall, and/or it might promote disease progression after PPE infusion by impairing protective SMC functions.

**Figure 1.**
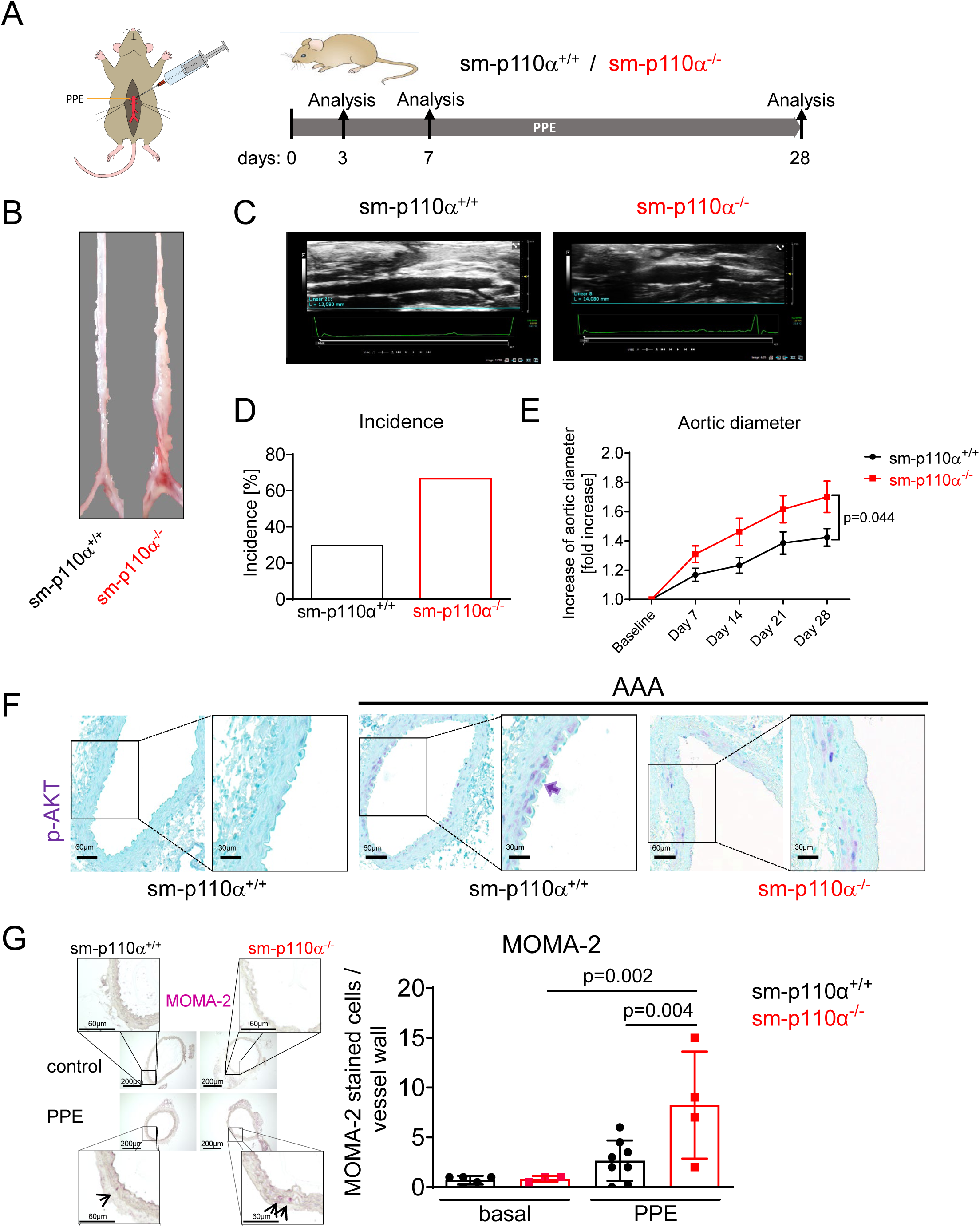
Lack of p110α in SMCs promotes AAA formation in mice. **A** Porcine pancreatic elastase (PPE) was infused into the infrarenal segment of aortae in male sm-p110α^+/+^ (n=10) and sm-p110α^−/−^ (n=9) mice. Aortae were analyzed under baseline conditions as well as 3, 7, and 28 days after PPE intervention. **B** Representative pictures of aortae and **C** ultrasound images from sm-p110α^+/+^ and sm-p110α^−/−^ mice demonstrate AAA formation 28 days after PPE infusion. **D** incidence for AAA (increase of lumen diameter >50%) (sm-p110α^+/+^: n=3 (10); sm-p110α^−/−^: 6 (9)). *P* value was calculated using Fisher’s exact test. **E** Analysis of ultrasound images of sm-p110α^−/−^ and sm-p110α^+/+^ mice revealed increase of the aortic diameter 0, 7, 14, 21 and 28 days after PPE intervention. Data is presented as means ± sem. *P* value was calculated mixed-effects analysis followed by Bonferroni correction. **F** Representative immunohistochemical staining showing AKT phosphorylation (purple) 3 days after PPE intervention in sm-p110α^−/−^ and sm-p110α^+/+^ mice compared to control. **G** MOMA-2 staining (red) of cross-sections from PPE and control sm-p110α^−/−^ and sm-p110α^+/+^ mice as indicated. Scale bars: 50 μm. Quantification on the right. n=3-7; *P* values were calculated using a 2-way ANOVA followed by uncorrected Fisher’s LSD. Data is presented as means ± S.D.

### p110α deficiency impairs aortic wall structure and function under basal conditions

We explored the impact of p110α deficiency in SMCs on aortic structure and function under basal conditions to identify potential preexisting vascular alterations. sm-p110α^−/−^ mice are characterized by a diminished aortic diameter compared to WT controls. The reduced vessel size likely reflects an impaired capacity of p110α^−/−^ SMCs to proliferate and migrate during aortic development (**Figure 2A**). Both, SMCs and ECM significantly contribute to aortic wall structure and function. Immunofluorescence staining for α-sm-actin and elastin as well as second harmonic generation to visualize collagen fibers did not show major defects in the aortic wall (**Figure 2B**). However, analysis of the ultrastructure by transmission electron microscopy revealed that abdominal segments of aortae from sm-p110α^−/−^ mice are characterized by derangements in the medial compartment, harboring detached SMCs, impaired elastic lamellae, and features of apoptotic cell death including fragmented nuclei and swollen mitochondria (**Figure 2C,D**). Next, we investigated the impact of p110α deficiency on the expression of phenotypic SMC marker genes. Bulk RNA-seq analysis revealed subtle but statistically significant downregulation of several contractile SMC marker genes (*Myh11*; *Smtn*; *Cnn1*; *Acta2*) in sm-p110α^−/−^ aortae compared to wild-type controls (**Figure 2E**). Western blot analysis demonstrated decreased protein expression of calponin and myosin heavy chain (MHC) in the wall of sm-p110α^−/−^ aortae, while the expression of α-sm-actin remained unchanged (**Figure 2F**). These findings might be related to both decreased SMC content and impaired SMC differentiation. Krüppel-like factor 4 (Klf4) is a master regulator of SMC de-differentiation and promotes the downregulation of contractile genes in SMCs.^28^ To test if p110α regulates calponin expression via Klf4 induction, we analyzed abdominal aortae under basal conditions. *Klf4* mRNA tended to be up-regulated in sm-p110α^−/−^ aortae compared with wild-type controls (**Figure 2G**). As expected, *Klf4* is induced by growth factors such as PDGF-BB in p110α^+/+^ SMCs, but this response remained unaltered in PDGF-BB treated p110α^−/−^ SMCs (**Figure 2H**), arguing against a role of Klf4 in p110α mediated regulation of contractile markers like calponin. Furthermore, increased mRNA expression of *CD68* and chemokine (C-C motif) ligand 2 (*Ccl2*) indicated elevated macrophage infiltration and inflammation in the sm-p110α^−/−^ abdominal segment (**Figure 2I, J**), but this may occur secondary to disturbances of the aortic wall structure under basal conditions.

**Figure 2.**
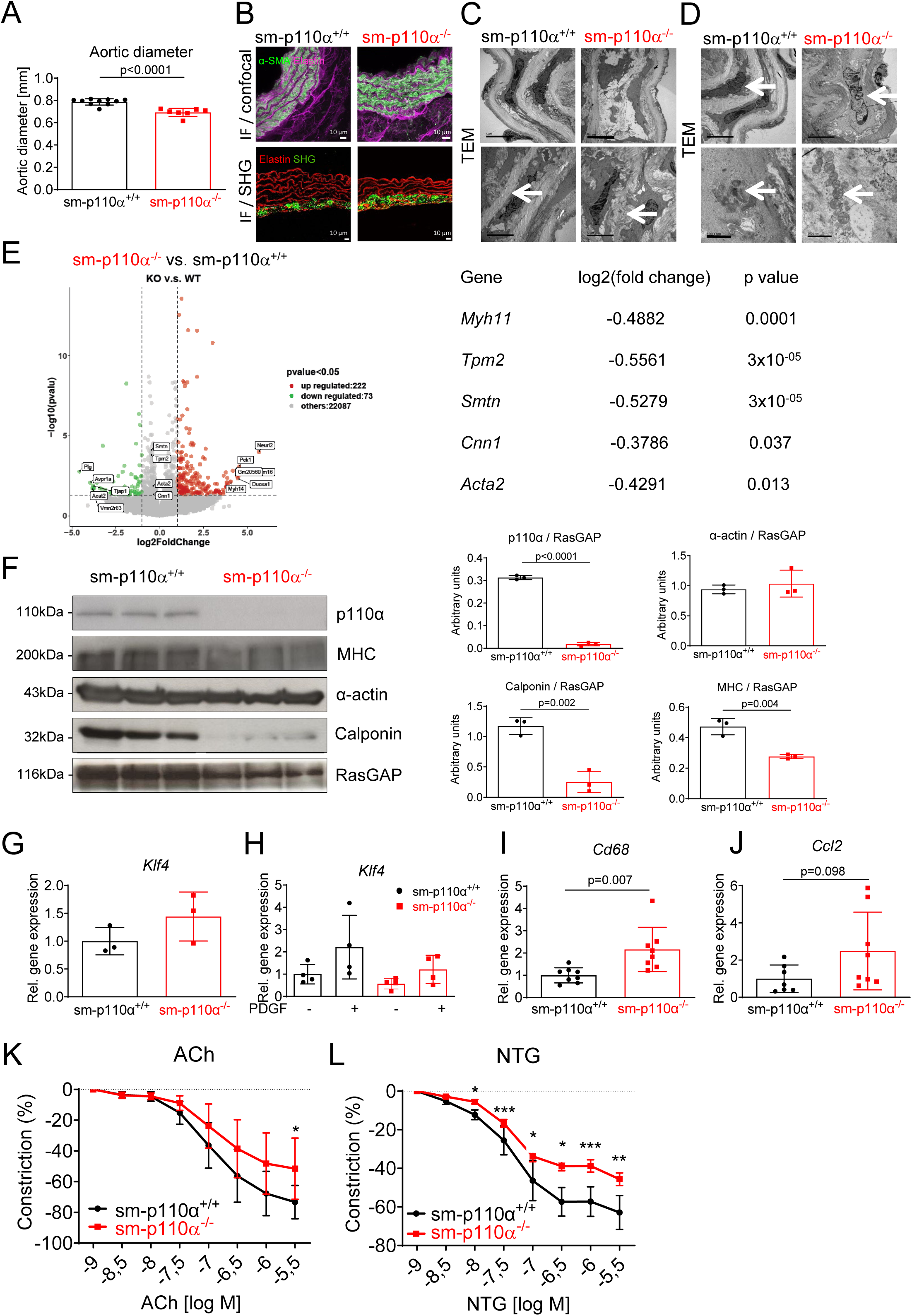
p110α deficiency impairs aortic wall structure and function under basal conditions. **A** Analysis of ultrasound images of infrarenal aortic segments from sm-p110α^−/−^ and sm-p110α^+/+^ mice under basal conditions and quantification of aortic diameter, *P* value was calculated using an unpaired two-tailed Student’s t-test. (n=7-10). **B** Representative images of immunofluorescence staining of α-sm-actin (green) and elastin (purple) (top) and second harmonic generation (SHG) (bottom) to visualize collagen (green) using confocal microscopy on aortic sections from sm-p110α^+/+^ versus sm-p110α^−/−^ mice. Scale bars: 50 μm. **C** Representative images of transmission electron microscopy (TEM) of abdominal aortas from sm-p110α^+/+^ and sm-p110α^−/−^ mice demonstrating an intact media structure (top) and intact elastic fibers (below, indicated by the white arrow) in sm-p110α^+/+^ mice (left) versus detached SMCs (top) and elastic fiber breaks (below, white arrow) in sm-p110α^−/−^ mice (right) (2500 x magnification). **D** Representative TEM-images demonstrating intact mitochondria (left, top) and nuclei (left, below) in sm-p110α^+/+^ mice versus disturbed mitochondria (right, top) and fragmented nuclei (right, below) in sm-p110α^−/−^ mice (12000 x magnification). Mitochondria and nuclei are indicated by white arrows, respectively. Scale bars: 5 μm. **E** Bulk RNAseq approach demonstrating differentially expressed SMC marker genes (right). Volcano plot on the left showing up- and downregulated genes in sm-p110α^+/+^ versus sm-p110α^−/−^ aortae (n=4, respectively) under basal conditions. *Myh* (myosin 11; *Tpm2*(tropomyosin b); *Smtn* (smoothelin); *Cnn1* (calponin 1); *Acta2* (α-sm-actin). (**F**) Western blot (left) and densitometric analysis (right) of protein expression of p110α and the differentiation markers α-sm-actin, calponin and myosin heavy chain (MHC) in sm-p110α^+/+^ and sm-p110α^−/−^ aortae (n=3, *P* value was calculated using an unpaired two-tailed Student’s t-test.) under basal conditions demonstrating impaired SMC differentiation in aortae lacking p110α in SMCs. RasGAP served as loading control. **G** mRNA expression of *Klf4* in sm-p110α^+/+^ and sm-p110α^−/−^ abdominal aortae under basal conditions (n=3). *P* value was calculated using an unpaired two-tailed Student’s t-test. **H** mRNA expression of *Klf4* in PDGF-BB (50 ng/ml) stimulated p110α^+/+^ and sm-p110α^−/−^ SMCs (n=3). *P* value was calculated using a 2-way ANOVA followed by uncorrected Fisher’s LSD. Data is presented as means ± S.D. **I** mRNA expression of macrophage marker *CD68* and (**J**) chemokine *Ccl2* in AAA from sm-p110α^+/+^ versus sm-p110α^−/−^ mice under basal conditions determined by quantitative PCR (n=7-8). *P* value was calculated using an unpaired two-tailed Student’s t-test. Data is presented as means ± S.D. **K** Relaxation of explanted abdominal aortic rings in response to acetylcholine (Ach) or (**L**) nitroglycerin (NTG) was assessed by isometric force measurements and expressed as % of maximal prostaglandin F2α-mediated constriction. *P* values were calculated using 2-way repeated measures ANOVA followed by Bonferroni correction. \**p* < 0.05, \*\**p* < 0.01. \*\*\**p*<0.001; n=4, respectively. Data is presented as means ± S.D.

To determine whether decreased expression of contractile marker genes and impaired aortic wall structure in knockout mice correlates with altered vascular function, we used wire myography to measure vasomotor properties of abdominal aortic specimens. Aortic rings isolated from sm-p110α^−/−^ mice displayed less relaxation in response to acetylcholine stimulation of endothelial nitric oxide (NO) production (**Figure 2K**). The dose response relationship in knockout aortae to the NO donor nitroglyceride was also altered (**Figure 2L**), indicating that this effect was endothelium independent and rather due to an impairment of SMC and aortic wall function.

Collectively, these data demonstrate preexisting structural and functional alterations in the aortic wall in sm-p110α^−/−^ mice. While these alterations alone did not promote AAA growth, data indicate that p110α signaling helps maintain the integrity of the aortic wall under physiological conditions. Thus, lack of p110α expression or activity may predispose to the development of AAA when additional triggers are present.

### Impact of p110α expression on vascular remodeling and SMC de-differentiation in AAA

We next aimed to identify disease-modifying properties of p110α signaling during AAA growth. AAAs are characterized by loss of SMCs in the media compartment. Regenerative capacities therefore depend on the ability of the aorta to compensate for cell loss. Analysis of histochemical staining revealed a sharply decreasing cellularity in the aortic wall 3 days after PPE treatment in the compromised segment (**Figure 3A,B**). Subsequently, the cellularity in the aortic wall gradually recovered, finally leading to increased media thickness after 28 days in both sm-p110α^+/+^ and sm-p110α^−/−^ mice (**Figure 3A-D**). However, this response was significantly reduced in sm-p110α^−/−^ mice (**Figure 3C,D**). PCNA staining indicated reduced proliferation of aortic cells in sm-p110α^−/−^ mice (**Figure 3E,F**), consistent with decreased cellular growth of aortic SMCs from sm-p110α^−/−^ mice *in vitro* (**Figure 3G**). These data suggest that p110α signaling contributes to SMC proliferation and thereby promotes vascular recovery and remodeling after PPE intervention.

**Figure 3.**
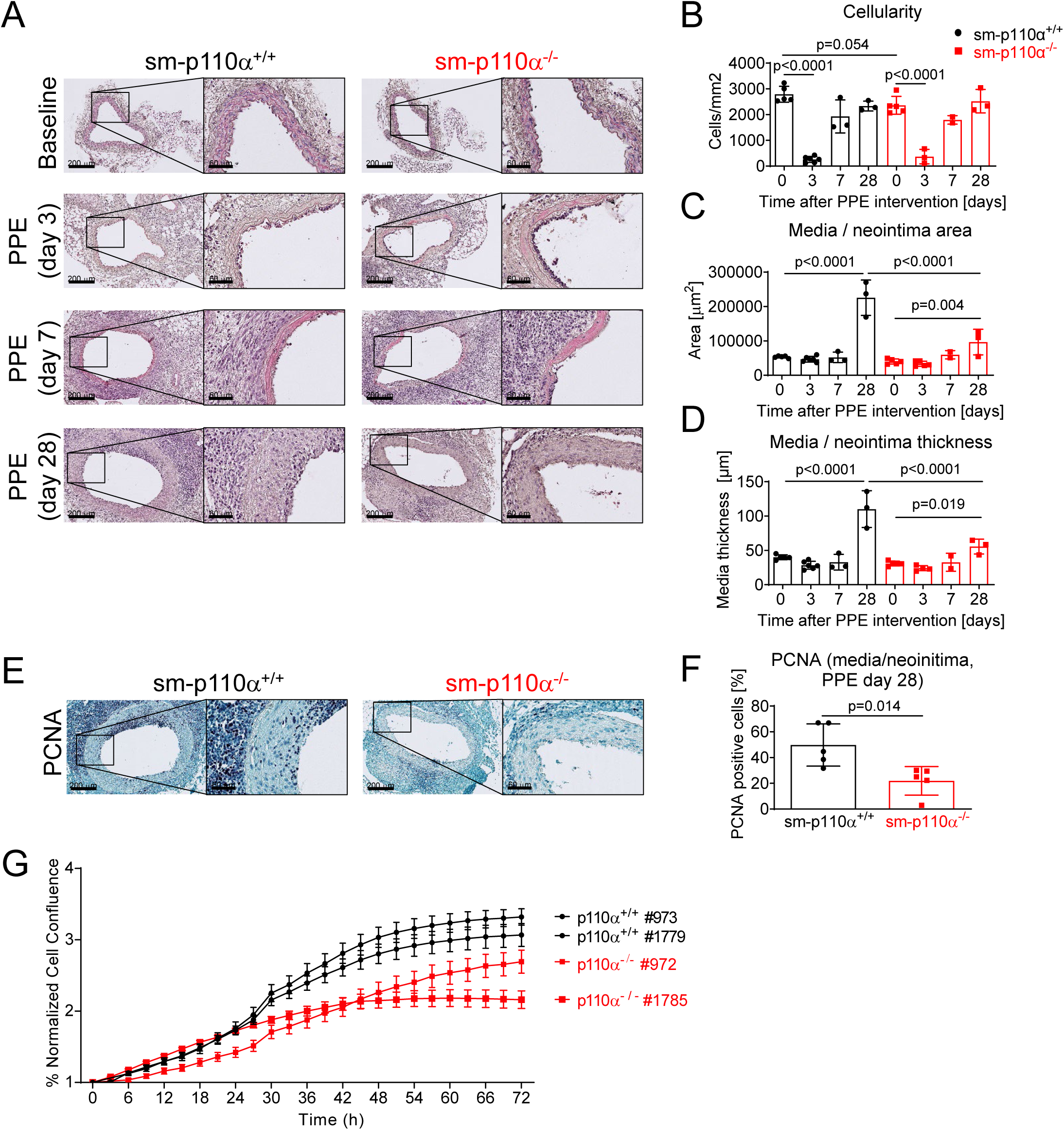
Impact of p110α expression on vascular remodeling and SMC differentiation in AAA. **A** Representative hematoxylin / eosin (H&E) stained cross sections of abdominal aortae from sm-p110α^+/+^ versus sm-p110α^−/−^ mice demonstrating vascular remodeling 0, 3, 7, and 28 days after PPE perfusion. Scale bars 200 μm (overview) and 60 μm (zoom). **B** Cellularity in the medial and neointimal compartment of indicated mice and periods of time. **C** Quantification of the media/neointima area and (**D**) analysis of media thickness (n=2-5 mice, respectively). *P* values were calculated using a 2-way ANOVA followed by Bonferroni correction. **E** Analysis of proliferating cells in the media/neointima compartment 28 days after PPE intervention in indicated mice. Sections from mice were stained for PCNA 28 days post PPE perfusion. Scale bars 200 μm (overview) and 60 μm (zoom). **F** PCNA+ cells were quantified in media and neointima compartments. n=5 mice. *P* value was calculated using an unpaired two-tailed Student’s t-test. Data is presented as mean ± S.D. **G** Analysis of SMC proliferation. Growing curves of aortic cells isolated from sm-p110α^+/+^ and sm-p110α^−/−^ mice (n=2, respectively). Data is presented as means ± SEM.

To determine the SMC content in AAA tissue, we analyzed the expression of the SMC differentiation markers calponin and α-sm-actin in aortic cross-sections from aortae before and 3, 7, and 28 days after PPE intervention by immunohistochemistry. Consistent with previous results, the percentage of the calponin-positive aortic wall area was significantly decreased in sm-p110α^−/−^ aortae under basal conditions. PPE intervention led to a pronounced decrease of calponin expression in both sm-p110α^+/+^ and sm-p110α^−/−^ aortae after three days and interestingly remained low until day seven, when cellularity had already increased (**Figure 4A,B**). This finding was confirmed by qRT-PCR (**Figure 4C**) and suggests that non-SMCs or fully de-differentiated SMCs must have migrated into the medial compartment prior to SMC reconstitution. Calponin protein expression was restored after 28 days, although this tended to be reduced in sm-p110α^−/−^ aortae (**Figure 4B**). Similar results were obtained with α-sm-actin immuno-staining and qRT-PCR (**Figure 4D-F**). However, consistent with Western blot analysis, α-sm-actin was not differentially expressed under basal conditions, whereas *Acta2* mRNA was decreased (**Figure 4F**). α-sm-actin protein expression was still reduced 28 days after PPE surgery, suggesting that SMC content and differentiation were not yet fully restored. Taken together, these findings indicate an initial loss of differentiated contractile SMCs after PPE intervention and a subsequent recovery phase, which is delayed in sm-p110α^−/−^ mice.

**Figure 4.**
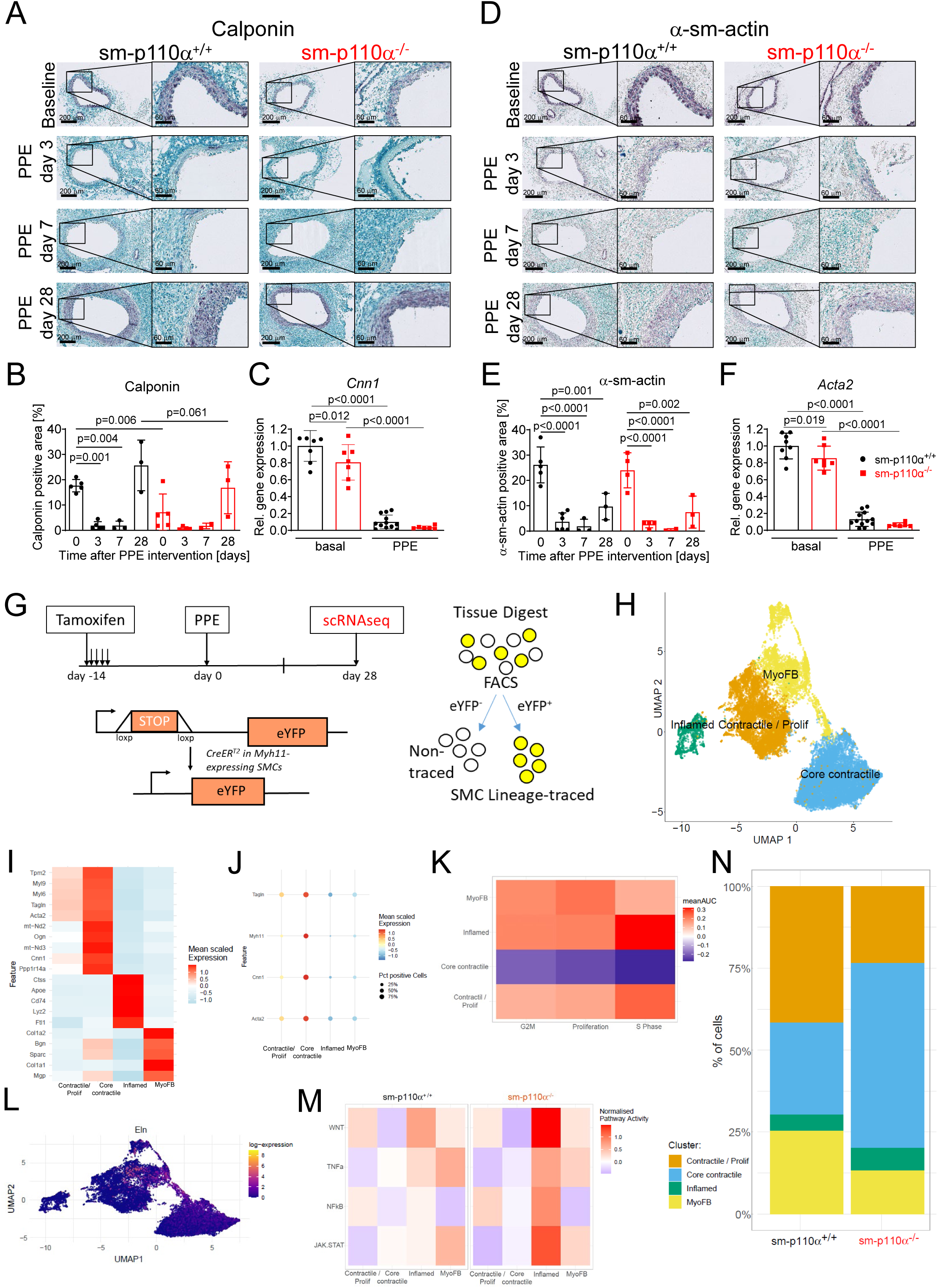
Effects of p110α deficiency on SMCs phenotypic modulation in PPE treated mice. Analysis of immunochemical staining and mRNA expression of calponin (**A-C**) and α-sm-actin (**D-F**) reveal time dependent changes after PPE surgery. Representative immunohistochemical staining of calponin (**A**) and α-sm-actin (**D**) expression in aortic cross-sections from sm-p110α^+/+^ and sm-p110α^−/−^ mice 0, 3, 7, and 28 days after PPE intervention, respectively (*n* = 3-5). Scale bars 200 μm (overview) and 60 μm (zoom). Quantification of protein expression is shown (**B**) Calponin and (**E**) (α-sm-actin). *P* values were calculated using a 2-way ANOVA followed by Bonferroni correction. mRNA expression of *Cnn1* (**C**) and *Acta2* (**F**) in sm-p110α^+/+^ and sm-p110α^−/−^abdominal aortae under basal conditions or 7 days after PPE intervention as indicated (n=6-12). *P* values were calculated using a 2-way ANOVA followed by uncorrected Fisher’s LSD. Data is presented as means ± S.D. **G** Cartoon depiction of SMC lineage tracing in ism-p110α^+/+^-eYFP and ism-p110α^−/−^-eYFP (Myh11-CreER^T2^ ROSA26-STOP^FL/FL^-eYFP / Pik3ca^fl/fl^) mice. Tamoxifen injection starting at day −14 induces Cre recombinase-mediated activation of eYFP in cells expressing myosin heavy chain 11 (*Myh11*) leading to permanent fluorescence in SMCs. Single-cell RNA sequencing (scRNA-seq) is performed 14 days after PPE intervention. **H** Uniform manifold approximation and projection (UMAP) of SMC sub-populations clusters (contractile/proliferative, core contractile, inflamed, myofibroblast-like). **I** Heatmap showing the expression of characteristic marker genes. **J** Dot plots show the expression of contraction markers (*Acta2, Cnn1, Myh11, Tagln*) in the SMC clusters. **K** Proliferation and cell cycle scores in the respective SMC clusters. **L** Feature plots demonstrate the expression of *Eln* in 4 SMC type clusters. Color scale indicates log2 expression values, from 0 (low) to 8 (high). **M** Heatmap of inflammatory pathway activity scores (Wnt, tumor necrosis factor (TNF), nuclear factor kappa B (NF-κB), Janus kinase-signal transducer, activator of transcription (JAK/STAT)) across SMC subpopulations in ism-p110α^+/+−^-eYFP versus ism-p110α^−/−^-eYFP. **N** Bar plots show cluster distribution within the SMC custers in ism-p110α^+/+−^-eYFP versus ism-p110α^−/−^-eYFP.

We next explored the effects of p110α deficiency on SMC de-differentiation in PPE treated mice. We generated tamoxifen-inducible SMC-specific conditional p110α KO mice with simultaneous SMC lineage tracing (Myh11-CreER^T2^ ROSA26-STOP^fl/fl^-eYFP *Pik3ca*^fl/fl^ or *Pik3ca*^WT/WT^ mice), referred to as ism-p110α^+/+^-eYFP and ism-p110α^−/−^-eYFP mice, respectively, after tamoxifen treatment. 14 days after PPE surgery, eYFP+ cells (SMCs and SMC-derived cells) from ism-p110α^+/+^-eYFP or ism-p110α^−/−^-eYFP AAA were subjected to scRNA-seq, since SMC related responses are expected to peak at this time point (own observation and ^15,29^) (**Figure 4G**). Uniform manifold approximation and projection (UMAP) analysis showed clusters of cells aligning with four distinct SMC sub-populations (contractile/proliferative, core contractile, inflamed, myofibroblast-like), based on the appropriate marker genes (**Figure 4H**). The specific cell marker expression characteristics are visualized in **Figure 4I**.

The contractile/proliferative and particularly the contractile SMC cluster highly expressed contraction-related genes such as *Myh11* and *Acta2*. The inflamed SMC cluster highly expressed immune-related genes such as *Ctss*, *Cd74* and *Lyz2,* and the myofibroblast-like SMC cluster extracellular-matrix-related genes such as *Col1a1/2* and *Bgn*. Expression analysis of contractile markers in the four subpopulations revealed that the core-contractile SMC subpopulation possessed the highest expression and the inflamed cluster the lowest (**Figure 4J**). In contrast, proliferation and cell cycle scores are enriched in contractile/proliferative, inflamed, and myofibroblast-like clusters, and decreased in the core contractile cluster, respectively (**Figure 4K**). Furthermore, whereas expression of contractile markers is more pronounced in core-contractile and contractile/proliferative SMC subgroups (**Supplementary Figure 2**), *Eln*, encoding tropoelastin, is enriched in myofibroblast-like SMCs, potentially indicating a reparative capacity of this subgroup in AAA (**Figure 4L**). Interestingly, pathway analysis revealed increased activation of inflammation related signaling pathways including Wnt, tumor necrosis factor (TNF), nuclear factor kappa B (NF-κB), Janus kinase-signal transducer and activator of transcription (JAK/STAT) pathways in inflamed SMCs from ism-p110α^−/−^-eYFP AAA (**Figure 4M**). Expectedly, the proportion of both, proliferating contractile/proliferative myofibroblast-like SMCs were lower in the ism-p110α^−/−^-eYFP sample compared with that in the control group (**Figure 4N**), indicating increased SMC proliferation of p110α expressing SMCs. In contrast, inflamed SMCs and core-contractile SMCs were increased in sm-p110α^−/−^ AAA. Reduced myofibroblast-like SMCs and increased inflamed SMCs in the ism-p110α^−/−^-eYFP group might indicate reduced reparative capacity and increased expression of ECM degrading enzymes in sm-p110α^−/−^ mice during AAA formation.

### p110α deficiency impairs elastic fiber homeostasis in AAA

ECM degradation was investigated in AAA from PPE treated sm-p110α^−/−^ and sm-p110α^+/+^ mice after different periods of time. Under basal conditions, p110α deficiency in SMCs caused significantly increased number of elastic fiber breaks in the thoracic segment (**Supplementary Figure 3A**) of the aorta, whereas the abdominal aorta was not affected (**Figure 5 A-C**). Analysis of fibrotic collagen content by Picro Sirius Red staining in aortae from sm-p110α^−/−^ mice and sm-p110α^+/+^ controls did not reveal substantial differences (**Supplementary Figure 3B,C**). After PPE surgery, destruction of the ECM peaked 28 days after PPE intervention in both wild-type controls and sm-p110α^−/−^ mice. However, elastic fiber degeneration occurred earlier in sm-p110α^−/−^ mice (**Figure 5A,B**), displaying significantly more elastic fiber breaks than control mice three days after PPE surgery (**Figure 5C**).

**Figure 5:**
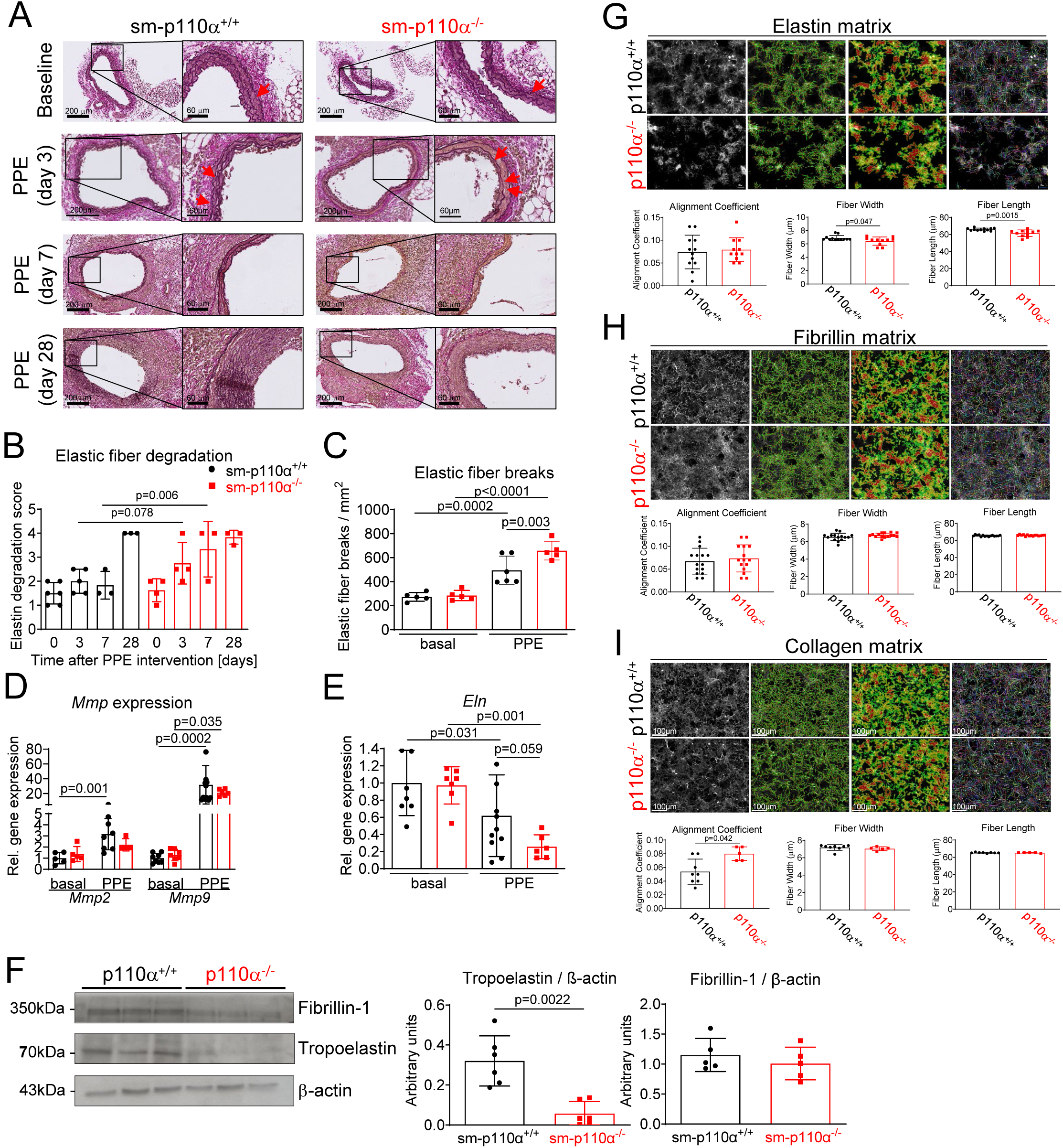
Elastic fiber homoeostasis in PPE treated sm-p110α^−/−^ mice and SMCs. **A** Representative images of elastica van Gieson staining in cross sections from sm-p110α^+/+^ versus sm-p110α^−/−^ mice either under basal conditions or 3, 7, and 28 days after PPE perfusion. Scale bars 200 μm (overview) and 60 μm (zoom). Elastic fiber breaks under basal conditions and at day 3 were indicated by red arrows. **B** Categorization of elastic fibers into score 1-5 (n=3-6). *P* values were calculated using a 2-way ANOVA followed by Bonferroni correction. **C** Quantification of elastic fiber breaks defined as interruptions in the elastic fibers, counted per cross-section, and shown in counts per mm^2^ in sm-p110α^+/+^ versus sm-p110α^−/−^ mice under basal conditions and 3 days after PPE perfusion, respectively. n=5-6. **D** mRNA expression of *Mmp2* and *Mmp9* and (**E**) *Eln* in AAA from sm-p110α^+/+^ versus sm-p110α^−/−^ mice 7 days after PPE perfusion determined by quantitative PCR (n=5-10). *P* values were calculated using a 2-way ANOVA followed by uncorrected Fisher’s LSD. Data is presented as means ± S.D. **F** Representative Western blots (left) and densitometrical analysis (right) demonstrating expression of elastin and fibrillin in SMCs from sm-p110α^+/+^ and sm-p110α^−/−^mice. RasGap and ß-actin served as loading controls. (n=5-6). *P* value was calculated using an unpaired two-tailed Student’s t-test. **G-I** Analysis of fiber width, length, and alignment coefficient. Aortic SMCs isolated from sm-p110α^+/+^ and sm-p110α^−/−^ mice were grown to confluence for seven days. Resultant elastin (**G**), fibrillin (**H**) and collagen (**I**) matrices were stained by immunofluorescence (representative images are shown in the first column (left), respectively). Scale bars: 50 μm. Fibrillar collagen alignment was quantified using CurveAlign/CT-FIRE. Second column: Images generated by CurveAlign showing the location (red dots) and orientation (green lines) of extracted fibers. Third row: Heatmaps indicating regional fiber alignment (red denotes areas of greater alignment). Fourth row: CT-FIRE extracted fibers overlaid on master image (first row). Information for entire fibers are shown and highlighted by different colored lines for contrast. Graphs illustrate mean fibrillar alignment, width, and length ± S.D. as indicated. Data normality was assessed using the Shapiro–Wilk test. Alignment coefficient data were analyzed by an unpaired, two-tailed Student’s T-test. Data in fiber width and fiber length were analyzed using the non-parametric Mann–Whitney U test, respectively. Data is presented as mean ± S.D.

During AAA formation, leukocyte and SMC derived matrix metalloproteinases (MMP)-2 and MMP-9 degrade ECM proteins and promote AAA formation. mRNA expression of *Mmp2* and *Mmp9* were significantly up-regulated in AAAs seven days after PPE intervention (**Figure 5D**). However, *Mmp2* and *Mmp9* were expressed at similar levels in both genotypes under basal conditions as well as after PPE intervention. Therefore, it is unlikely that MMP-2 and MMP-9 derived from leukocytes or inflamed SMCs contributed to increased elastic fiber breaks in the knockout animals at this time-point. In contrast, seven days after PPE infusion, mRNA expression of tropoelastin (*Eln*) was diminished in sm-p110α^−/−^ mice compared to wild-type (**Figure 5E**). This prompted us to explore the impact of p110α deficiency on the expression of elastic fiber components in SMCs *in vitro*. p110α^−/−^ SMCs produced less tropoelastin than wild-type control cells, whereas fibrillin-1 expression was unaffected (**Figure 5F**). Further matrix analysis of immuno-stained ECM fibers in SMC cultures elucidated the effects of altered ECM expression on the respective fiber structure. Quantitative analysis of alignment, fiber width, and fiber length revealed that elastin fibers from p110α deficient SMCs are significantly shorter and thinner than those from control cells, whereas fibrillin fiber dimensions were not compromised (**Figure 5G,H**). These data indicate that disturbed elastin homeostasis in p110α^−/−^ SMCs might be a prerequisite for increased elastic fiber breaks in sm-p110α^−/−^ aortae. In addition, we found a significantly increased alignment coefficient of collagen fibers in p110α^−/−^ SMC cultures (**Figure 5I**), indicating that p110α^−/−^ deficiency in SMCs promoted a more organized collagen structure.

### Inactivation of FOXO1 rescues the sm-p110α^−/−^ phenotype and protects against aortic aneurysm formation

To elucidate relevant p110α mediated signaling events in more detail, we investigated growth factor stimulated aortic SMCs from sm-p110α^−/−^ and sm-p110α^+/+^ mice. PDGF-BB stimulation of p110α expressing wild-type SMCs, rapidly induced phosphorylation of numerous signaling molecules downstream of PI3K such as AKT (including isoforms AKT1 and AKT2), FOXO transcription factors (FOXO1, FOXO3, FOXO4), and glycogen synthase kinase 3β (GSK3β). These responses were significantly reduced in p110α^−/−^ SMCs (**Figure 6A**), while total protein expression was unaltered (**Supplementary Figure 4A)**. Application of insulin led to similar results (**Supplementary Figure 4B**). It was previously demonstrated that growth factor mediated inactivation of FOXO1 is implicated in SMC proliferation.^30^ Consequently, we analyzed the impact of FOXO1 inhibition on PDGF-BB stimulated proliferation of p110α^−/−^ SMCs. p110α deficiency significantly diminished SMC proliferation compared to wild-type cells. Strikingly, the restricted mitogenic response of p110α^−/−^ SMCs was fully rescued by application of the specific FOXO1 inhibitor AS1842856, whereas FOXO1 inhibition had no effect on PDGF-BB stimulated wild-type cells (**Figure 6B**). These data demonstrate that p110α mediated inactivation of FOXO1 is mechanistically responsible for PDGF-BB induced SMC proliferation. Furthermore, siRNA mediated knockdown of *Foxo1* increased *Eln* as well as *Cnn1* expression in SMCs, suggesting that FOXO1 inactivation might be important for phenotypic modulation and elastogenesis (**Figure 6C**). To analyze the impact of FOXO1 inactivation on AAA formation, we generated smooth muscle specific FOXO1 deficient sm-FOXO1^−/−^ mice and subjected them to the PPE mouse model (**Figure 6D**). Ultrasound examination 28 days after PPE intervention (**Figure 6E**) revealed decreased incidence (67% versus 40%) of AAA disease and a significantly reduced increase in aortic diameter in sm-FOXO1^−/−^ mice, respectively (**Figure 6F,G**).

**Figure 6.**
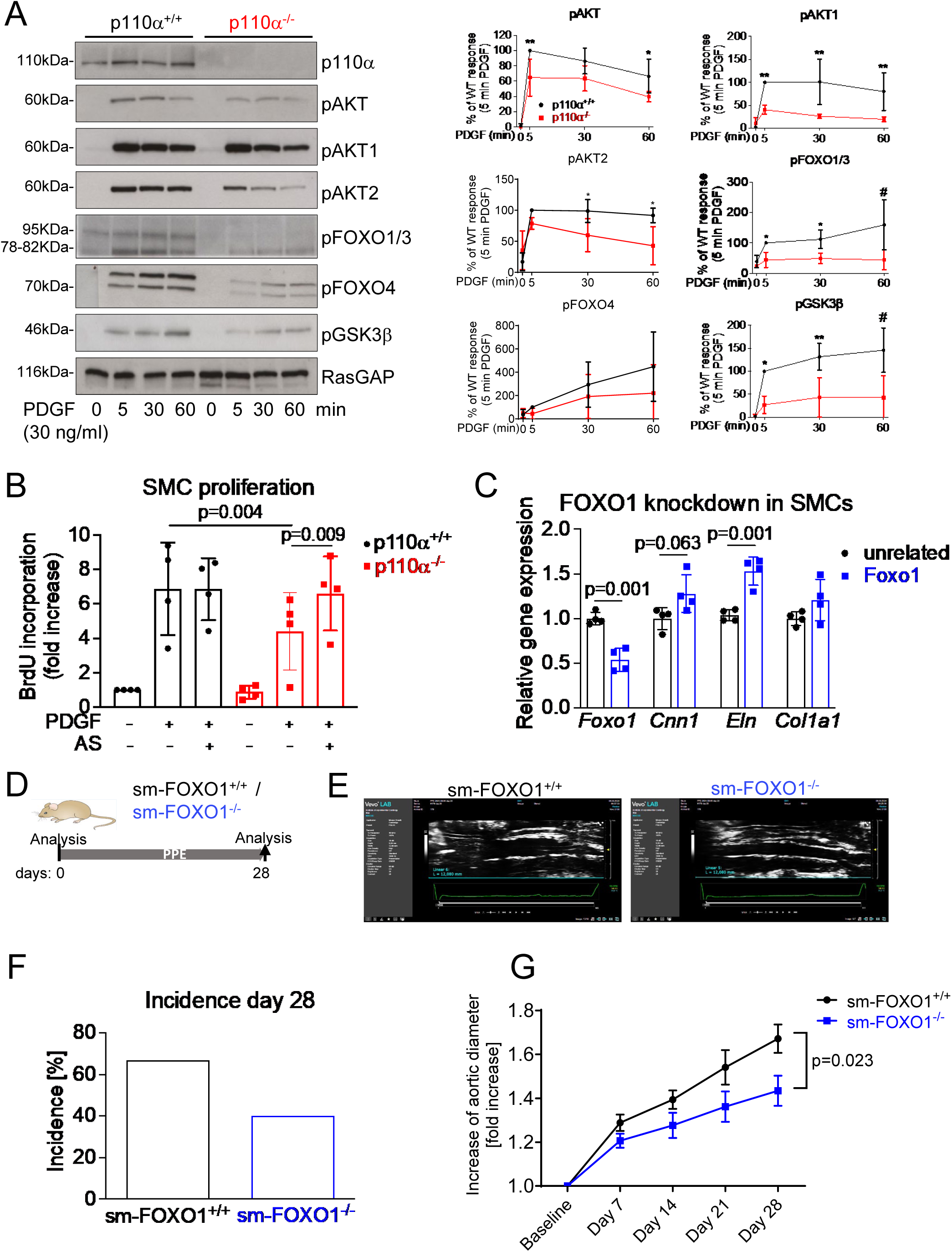
Inactivation of FOXO1 in SMCs rescues the p110α^−/−^ phenotype and protects against AAA formation. A Representative western blots of cultured PDGF-BB (50 ng/ml, 0-60 min) stimulated SMCs showing phosphorylated signaling proteins downstream of p110α as indicated. Densitometrical analyses are shown on the right. (n=3-4). *P* values were calculated using a 2-way ANOVA followed by Bonferroni correction. \**p* < 0.05, \*\**p* < 0.01. #*p*<0.001. **B** PDGF-BB (50ng/ml) dependent SMC proliferation in p110α^−/−^ SMCs and WT controls in the absence or presence of the FOXO1 specific inhibitor AS1842856 (AS, 1µM). n=5. *P* values were calculated using a 2-way ANOVA followed by Bonferroni correction. **C** mRNA expression of *Foxo1*, *Cnn1*, *Eln*, and *Col1a1* in SMCs after knockdown of *Foxo1* by four different siRNAs or in the presence of four different unrelated siRNAs as control determined by quantitative PCR. *P* value was calculated using an unpaired two-tailed Student’s t-test. **D** Aortae of male sm-FOXO1^+/+^ (n=9) and sm-FOXO1^−/−^ (n=10) mice were infused with PPE and analyzed before and 28 days after the intervention. **E** Representative ultrasound pictures of aortae from sm-FOXO1^+/+^ and sm-FOXO^−/−^ mice demonstrate AAA formation 28 days after PPE infusion. **F** Incidence for AAA (increase of lumen diameter >50%, (sm-FOXO1^−/−^: n=6 (9); sm-FOXO1^+/+^: n=4 (10)). *P* value was calculated using Fisher’s exact test. **G** Analysis of ultrasound images of sm-FOXO1^−/−^ and sm-FOXO1^+/+^ mice revealed increase of the aortic diameter 0, 7, 14, 21 and 28 days after PPE intervention. *P* value was calculated mixed-effects analysis followed by Bonferroni correction.

Taken together, inactivation of FOXO1 rescues impaired proliferation of sm-p110α^−/−^ SMCs, promotes calponin and elastin expression, and protects against AAA formation.

### Pharmacological activation of p110α protects against aortic aneurysm formation

To test p110α signaling as a pharmacological target to treat AAA disease, we used the recently developed p110α isoform selective activator UCL-TRO-1938.^24^ Accordingly, UCL-TRO-1938 activated AKT signaling and cell proliferation in wild type SMCs, whereas it was completely ineffective in p110α^−/−^ SMCs (**Figure 7A,B**). Furthermore, UCL-TRO-1938 induced proliferation was significantly diminished in FOXO1^−/−^ SMCs, again demonstrating the importance of FOXO1 inactivation in p110α dependent mitogenic signaling (**Figure 7C**). To assess the effects of UCL-TRO-1938 on AAA formation, we continuously applied the activator via osmotic mini-pumps (0.11 mg/kg/d) starting three days before PPE surgery until day 28 (**Figure 7D**). Analysis of ultrasound images revealed that pharmacological activation of p110α protected against AAA formation in C57Bl/6 wild type mice (**Figure 7E,F**). Importantly, this protective effect was diminished in sm-FOXO1^−/−^ mice (**Figure 7G,H**), whereas UCL-TRO-1938 ameliorated AAA progression in sm-FOXO1^+/+^ control mice (**Figure 7I,J**). These findings suggest that UCL-TRO-1938 mainly exerted its protective effects via p110α dependent FOXO1 inactivation in SMCs. To explore a potential relevance of activating growth factor-RTK-p110α-AKT-FOXO1 signaling in human AAA disease, we determined tissue samples from AAA patients. We found significant mRNA upregulation of *PDGFB* and *FOXO1*, while gene expression of *PIK3CA* (p110α) itself was not significantly altered (**Figure 7K-M, Supplementary Table 1**). Together, our findings suggest that FOXO1 in SMCs promotes AAA formation, which is consistent with its known pro-inflammatory effects in macrophages in aortic disease^31^, and that pharmacological p110α activation exerts protective effects via inactivation of FOXO1, which is specifically upregulated in human AAA.

**Figure 7.**
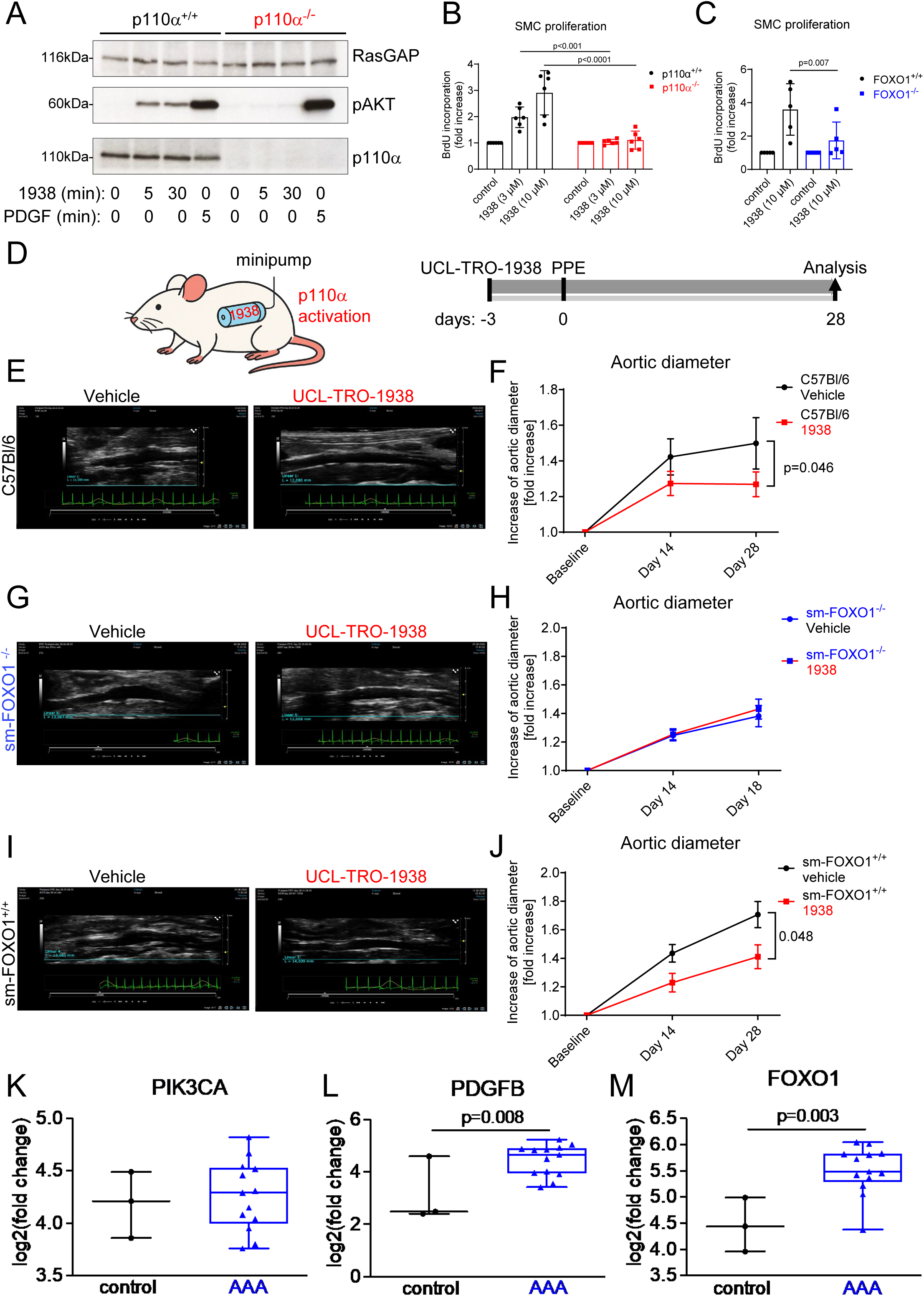
Pharmacological activation of p110α protects against aortic aneurysm formation. **A** Representative western blots of cultured PDGF-BB (50 ng/ml) versus UCL-TRO-1938 (10µM) stimulated SMCs showing AKT phosphorylation and protein expression of p110α. RasGAP served as loading control. **B** UCL-TRO-1938 stimulated SMC proliferation (BrdU incorporation) in p110α^+/+^ and p110α^−/−^ cells (n=6) and **C** in FOXO1^+/+^ and FOXO1^−/−^ cells (n=5). *P* values were calculated using a 2-way ANOVA followed by uncorrected Fisher’s LSD. **D** ULC-TRO-1938 (1mM) loaded osmotic minipumps (Alzet, 2004) were subcutaneously implanted 3 days prior to PPE intervention in C57BL/6J (**E, F**), sm-FOXO1^−/−^ (**G, H**), sm-FOXO1^−/−^, and (**I, J**) sm-FOXO1^+/+^ mice. **E, G, I** Representative ultrasound images of aortae from vehicle and UCL-TRO-1938 treated C57BL/6J (**E**), sm-FOXO1^−/−^ (**G**), and (**I**) sm-FOXO1^+/+^ mice, respectively, demonstrate AAA formation 28 days after PPE infusion. **F, H, J** Analysis of ultrasound images of vehicle and UCL-TRO-1938 treated mice (C57BL/6J(n=8-9) (**F**), sm-FOXO1^−/−^ (n=5-7) (**H**), sm-FOXO1^+/+^ (n=4-5) (**J**)) revealed increase of the aortic diameter 14 and 28 days after PPE infusion. *P* value was calculated by mixed-effects analysis followed by Bonferroni correction. **K, L, M** mRNA expression of *PIK3CA* (p110α) (**I**) *PDGFB* (platelet-derived growth factor B-chain) (**J**), and *FOXO1* (**K**) in human aortic samples from healthy controls (n=3) or AAA patients (n=12). *P* value was calculated using an unpaired two-tailed Student’s t-test.

## Discussion

The findings reported herein indicate that loss of p110α signaling in SMCs results in impaired SMC functionality, contributing to disturbed aortic wall structure and function as well as exaggerated development of AAA in PPE treated mice. Consequently, pharmacological activation of p110α ameliorates AAA disease (**Graphical abstract**). Thus, our data suggest that p110α signaling – via FOXO1 inactivation – exerts relevant protective functions in the aortic wall under physiological conditions as well as in AAA disease, and its activation arises as a potential therapeutic approach.

These findings should be considered alongside our previous work demonstrating that SMC-specific depletion of p110α attenuates vascular remodeling in other vascular diseases, including neointima formation after balloon injury or pulmonary hypertension.^18,19^ However, these diseases are characterized by excessive SMC proliferation and hyperplasia of the vessel wall, whereas aneurysms are characterized by loss of SMCs, suggesting that p110α exerts context-dependent effects on balancing vascular homeostasis.

The aorta of AAA patients is characterized by a decreased number of SMCs compared to healthy vessels,^32,33^ and the frequent appearance of apoptotic SMCs,^34,35^ leading to thinning of the aortic wall.^5^ Reduced SMC numbers impair connective tissue production and maintenance of elastin fiber integrity, promoting ECM degradation, inflammation, and AAA formation.^33^ Consistent with this situation in human AAAs, sm-p110α^−/−^ aortae displayed reduced wall thickness and features of apoptotic cell death including fragmented nuclei and swollen mitochondria. This was expected since p110α mediates SMC survival, and lack of p110α promotes SMC death.^18,19,36^ Additionally, sm-p110α^−/−^ mice exhibited disturbances in the aortic wall structure, reduced expression of contractile genes (calponin, MHC), and decreased responsiveness to the vasodilator NO under basal conditions. These impairments suggest that p110α plays an important role in the development and maintenance of a functional aortic wall. It is likely that the subtle pre-existing vascular derangements, which did not result in an obvious phenotype under basal conditions, make sm-p110α^−/−^ aortae susceptible to a second AAA promoting trigger. As a result, sm-p110α^−/−^ mice displayed significantly larger aneurysms compared to wild-type, using the PPE model. This model is considered to best mimic human AAA disease^37,38^, leading to time-dependent pathogenesis with acute to chronic inflammation and vascular wall remodeling,^29,39^ resembling end stage human AAA disease.^40,41^

Our data implicate that p110α is crucial for maintaining vascular integrity, but also controls regenerative mechanisms that counteract aortic dilatation during AAA formation. To regenerate the compromised vessel segment, loss of SMCs must be compensated by increased cell proliferation or infiltration. Indeed, diminished proliferative capacity is seen in human AAA-derived versus non-aneurysmal SMCs.^9^ We showed that p110α signaling in SMCs contributed to increased proliferation of medial cells, increased aortic wall area, and a trend toward increased percentage of Calponin+ SMCs 28 days after PPE injury, together indicating that p110α activation elevated SMC numbers in the media. Nevertheless, the impact of SMC proliferation on AAA progression remains controversial. For example, mTOR-dependent proliferative SMCs promote dilatation and dissection of the aortic wall rather than prevent disease progression,^42^ and inhibition of tyrosine kinase mediated SMC proliferation and phenotypic switching by lenvatinib ameliorates AAA formation.^43^ On the other hand, long non-coding RNA NUDT6 mediated SMC proliferation and survival protect against AAA disease.^44^ These contradictory findings may be explained by the fact that SMC proliferation is tightly coupled to SMC de-differentiation, both having distinct effects on AAA disease. Following vessel injury, SMCs de-differentiate and undergo clonal expansion as demonstrated in angiotensin II–induced thoracic aortic aneurysm or PPE induced AAA.^15,17^ SMCs can de-differentiate into distinct SMC subtypes, which might exert opposing functions in AAA disease.^12–14,28,42^

To investigate the impact of p110α deficiency on SMC cluster composition during AAA development, we lineage-traced eYFP+ SMCs and SMC derived cells from ism-p110α^+/+^-eYFP and ism-p110α^−/−^-eYFP mice, 14 days after PPE surgery, when SMC repopulation occurred in the compromised aortic segment (own observation and ^15^). Similar to what was previously demonstrated in an elastase-induced AAA model,^15^ we identified four distinct SMC clusters. Core-contractile SMCs displayed highest expression of contractile marker genes and lowest expression of genes involved in cell proliferation, therefore most likely representing quiescent contractile SMCs. Cell cycle scores were elevated in three proliferative SMC clusters including contractile/proliferative and myofibroblast-like SMCs, which were decreased in ism-p110α^−/−^-eYFP AAA, most probably due to the lack of p110α mediated mitogenic effects. Despite restricted PI3K mediated proliferation, we observed an increased proportion of inflamed SMCs in ism-p110α^−/−^-eYFP mice displaying increased activation of inflammatory pathways including Wnt, TNF, NF-κB, and JAK/STAT, which are implicated in cytokine and MMP production and promote AAA progression.^45^ These findings suggest that p110α signaling attenuated SMC de-differentiation towards an inflamed phenotype.

Recent studies demonstrated that loss of proliferative/contractile SMCs or myofibroblast-like SMCs in aortic tissues were associated with aortic dissection and aneurysm in mice, suggesting that these SMC subtypes contribute to aortic repair, remodeling, and reinforcement of the vessel wall by maintaining vascular homeostasis and compensating for SMC loss.^16,46–48^ In addition, inflamed SMCs facilitate AAA progression by secreting inflammatory chemokines and promoting MMP production,^49,50^ particularly MMP-2 and MMP-9, which directly degrade collagen and elastin.^51^ Altered SMC cluster distribution in mice lacking p110α in SMCs might therefore be linked to both impaired aortic tissue regeneration and increased elastic fiber degradation as shown in sm-p110α^−/−^ mice. In accordance with a reduced content of myofibroblast-like SMCs and increased inflamed SMCs in ism-p110α^−/−^-eYFP mice, we found a strong trend towards decreased expression of *Eln* and disturbed elastin-fiber homoeostasis in sm-p110α^−/−^ mice. However, we did not observe increased *Mmp2* and *Mmp9* expression, respectively, seemingly arguing against a prominent role of inflamed SMCs in MMP mediated elastic fiber degradation. Notably, scRNA-seq data were collected two weeks after PPE surgery, whereas the expression of MMPs in AAA from sm-p110α^−/−^ mice was analyzed after seven days at the peak of the inflammatory response. Therefore, we cannot exclude an impact of inflamed SMCs, particularly in the later stages of AAA progression. Furthermore, we detected increased activation of inflammation related signaling pathways in inflamed p110α^−/−^ SMCs which might have modulated and enhanced immune responses in AAA, thereby aggravating the disease. This would be consistent with data relating to an increased content of MOMA2^+^ cells in sm-p110α^−/−^ AAA at day 28. Collectively, these findings suggest that p110α exerted protective effects during AAA progression by elevating both, aortic SMC content and regenerative capacities, and potentially by mitigating SMC inflammation.

We identified inactivation of the transcription factor FOXO1 as the crucial downstream event for p110α mediated SMC proliferation, as specific inhibition of FOXO1 completely rescued the anti-proliferative p110α^−/−^ phenotype. Furthermore, whereas pharmacological p110α activation ameliorates AAA disease in wild type animals, it was ineffective in sm-FOXO1^−/−^ mice. We show that p110α activation promotes phosphorylation of FOXO1 without altering its protein expression. Numerous studies demonstrated that PI3K/AKT dependent phosphorylation of FOXO1 results in nuclear exclusion of FOXO1 attenuating its transcriptional activity.^52^ These findings collectively support the conclusion that inactivation of FOXO1 downstream of p110α is sufficient to protect against AAA progression. Since FOXO1 deficiency in smooth muscle cells protected against AAA progression, up-regulation of FOXO1 in human AAA indicates a possible disease-promoting role in patients and thus underscores the therapeutic potential of pharmacological inhibition of FOXO1 or activation of p110α. However, its actions in SMCs are not well understood thus far. It is suggested that PI3K/AKT dependent phosphorylation of FOXO1 mediates downregulation of the cell cycle inhibitor p27^kip^, thereby promoting cellular proliferation.^30^ In addition, it was recently shown that insulin dependent activation of PI3K/AKT/FOXO1 signaling prevents SMC inflammation in atherosclerosis via downregulation of thrombospondin.^53^ This finding indicates that PI3K dependent inactivation of FOXO1 attenuates de-differentiation towards inflamed SMCs and is consistent with our own data showing increased content of inflamed SMCs in ism-p110α^−/−^-eYFP AAA. However, recent data reveal that conditional knockout of thrombospondin in SMCs did not affect aortic dissection in the murine AA-Ang II-model.^54^ Hence, it remains elusive whether FOXO1 mediated SMC inflammation contributes to AAA progression.

Our data reveal beneficial effects of the p110α-activating compound UCL-TRO-1938 on AAA disease in WT mice but not in mice lacking FOXO1 in SMCs. These findings demonstrate that (a) p110α is a drugable target to treat AAA disease; (b) enhanced p110α signaling exerts protective functions during AAA progression; and (c) FOXO1 in SMCs is a primary target of pharmacological p110α activation. A recent study suggests that UCL-TRO-1938 and RTKs synergistically activate p110α.^24^ In this context it is important to note that we found *PDGFB* upregulated in human AAA patients suggesting increased PDGF receptor activity. These findings suggest a potential of p110α-activation for use in tissue protection and regeneration in AAA disease. However, since gain of function mutations in the *PIK3CA* gene are frequently observed in human cancers, the question arises, whether the pharmacological activation of p110α could potentially exert oncogenic effects. Notably, mutant *PIK3CA* alone is a weak driver oncogene. Mice constitutively expressing the *Pik3ca^H1047R^*hot-spot mutation do not develop cancer within a year.^24,55^ Furthermore, pharmacological activation is different from constitutive p110α activation. Whereas UCL-TRO-1938-induced p110α signaling is attenuated upon prolonged exposure by endogenous feedback mechanisms, oncogenic *PIK3CA* activation causes constitutive uncontrolled p110α signaling.^24^ Although long-term oncogenic risk has not been tested here, these data suggest that UCL-TRO-1938 mediated p110α activation is unlikely to cause or promote cancer.^24^

In conclusion, our data demonstrate that p110α signaling promotes vascular integrity and regeneration in healthy and diseased aorta by maintaining contractility, elastogenesis, and proliferative capacities of aortic SMCs. Promotion of p110α/FOXO1 signaling allows to specifically induce SMC proliferation without promoting detrimental de-differentiation towards inflamed SMCs. This is of particular importance to prevent or treat even late stage AAAs. Since a specific p110α activator and a FOXO1 inhibitor are readily available^24,56^, the findings presented here may pave the way for the development of new therapeutic or preventive strategies against AAA disease. Moreover, the impact of p110α/FOXO1 signaling on vascular integrity may potentially be clinically relevant in the context of current approaches aimed at chronic p110α inhibition for cancer therapy.

## Acknowledgements

The technical support by Frank Oberhäuser, Maximilian Becker, Nadja Klein, and Simon Grimm is greatly acknowledged. This works contains part of a doctoral thesis by MS. We thank Beatrix Martiny and Christian Jüngst (CECAD Imaging Core Facility, University of Cologne) as well as Lena Schumacher at the FACS & Imaging Core Facility at the Max Planck Institute for Biology of Aging (University of Cologne, Cologne, Germany) for excellent technical support.

