## Supplementary Material for "PI 3-kinase isoform p110α mediated FOXO1 inactivation protects against aortic aneurysm formation"

**Material and Methods** 1

UCL-TRO-1938 application via osmotic minipumps 1
Isometric force measurements 1
Bulk RNAseq 1
Generation of ism-p110α$^{+/+}$-eYFP and ism-p110α$^{-/-}$-eYFP lineage-tracing mice 2
Single Cell Sequencing - Cell processing for single-cell RNA sequencing 2
Single Cell Sequencing - Cell Isolation 3
Single Cell Sequencing - Transcriptome Preprocessing 3
Single Cell Sequencing - Data Processing and Analysis 3
Cell Culture and treatment with siRNAs 3
In vitro elastic fiber matrix analysis 4
Proliferation assay 4
BrdU incorporation 4
Immunoblotting 5
Immunohistochemistry & histochemistry 5
Second harmonic generation 6
Real-time qPCR 6
Materials and antibodies 6

**References** 7

**Supplementary Figure 1:** Increase of the aortic diameter in mm. 8

**Supplementary Figure 2:** Feature plots with expression of contractile genes. 9

**Supplementary Figure 3:** Elastic fiber breaks and collagen expression. 10

**Supplementary Figure 4:** p110α signaling in PDGF and insulin stimulated SMCs. 11

**Supplementary Figure 5:** Gating strategy to isolate YFP+ cells. 12

**Supplementary Table 1:** Patient Information. 13

### Material and Methods

#### UCL-TRO-1938 application via osmotic minipumps

To investigate the effect of p110α activation on AAA development, C57Bl/6J mice undergoing the PPE-induced AAA model were implanted with UCL-TRO-1938 loaded osmotic minipumps (Alzet model 2004) 3 days prior to PPE intervention. For implantation, the animals were anesthetized by inhalation of isoflurane (5% initial, 2% maintenance) and analgesed by subcutaneous injection of carprofen at 5 mg/kg body weight, 30 minutes before the start of surgery. The animals were secured on a heating pad to protect them from cooling down. Furthermore, they received Eye Care Gel (Remend Cornea®) to protect the eyes from drying out. Subsequently, a skin incision approximately 10 mm in length was made in the neck region, and a pocket was formed (approx. 6 cm), in which the pump was placed subcutaneously. The wound was closed with 3 wound clips (Reflex, 7 mm). UCL-TRO-1938 (1mM) was dissolved in 10% DMSO: 90% methylcellulose (0.5% w/v MC plus 0.1% TWEEN-80 v/v)) and administered at a rate of 0.25 µl/hr resulting in a daily dose of approximately 0.11 mg/kg body weight per day. Controls received pumps filled with the solvent.

#### Isometric force measurements

Isometric force measurements were performed as described.[1] Briefly, mice were anaesthetised using isoflurane (3–4%) in an anaesthesia chamber, transferred to a heating pad for inhalation anaesthesia, and kept under anaesthesia using an anaesthesia mask (7% isoflurane). The animals receive a subcutaneous injection of carprofen (5 mg/kg body weight) for analgesia 30 minutes before the procedure. After thoracotomy, explanted sections of the aortas were immediately kept in Krebs-Henseleit buffer on ice. 2-mm segments were mounted onto steel wires connected to force transducers of a standard organ bath filled with 25 ml Krebs-Henseleit buffer at 37°C and gassed with carbogen. Each ring was equilibrated for 30 minutes and continuously stretched up to a tension of 1.1 g. KCl (80 mM) was added twice to each bath with subsequent washing. To create a dose-response curve to PGF2α, we added it to the bath with concentration (in µM) of 0.1, 0.2, 0.4, 1, 2, 4, and 10. Thereafter, a dose responses to acetylcholine (Ach) or nitroglycerine (NTG) were recorded with concentrations from $1 \times 10^{-9}$ M in steps of half log10 levels up to a concentration of $1 \times 10^{-5.5}$ M.

#### Bulk RNAseq

After total RNA was extracted, RNA purity and integrity were checked by microfluidic electrophoresis using the Bioanalyzer (Qiagen, Hilden Germany). Following the quality control procedure, mRNA was enriched using Oligo(dT) magnetic beads and randomly fragmented for cDNA synthesis as previously described. Libraries were generated using the NEBNext UltraTM RNA Library Prep Kit (New England Biolabs, Ipswich, MA, USA) for the Illumina system following the manufacturer’s instructions. After cluster generation, single-end sequencing was performed using the Illumina HiSeq 2500 platform. Adapters were trimmed from raw reads, which were subsequently filtered by removing reads with a low quality score (50% bases of the read is <= 5). To accomplish the alignment, read mapping to the hg18 reference genome was carried out by means of STAR. Then htseq-count was used to quantity mapped reads counts for every gene of each sample. Number of mapped reads was calculated and gene expression levels were presented as FPKM (Fragments Per Kilo bases per Million reads). Subsequently, the differential expression and sample distance of two conditions was calculated using the DESeq2 R package. And top 100 differentially expressed genes (DEGs) ordered by p-value were selected to reflect relative expression levels across samples using pheatmap.

RNA-seq profiles were generated from 16 human samples (3 healthy samples and 13 diseased samples) and 8 murine samples (4 sm-p110α$^{+/+}$ samples and 4 sm-p110α$^{-/-}$ samples). Thus, in total 16 human samples and 8 murine samples were assessed for transcriptomic changes. DEGs were identified using a statistical threshold of $P < 0.05$ and fold change $\geq 2$.

**Generation of ism-p110α$^{+/+}$-eYFP and ism-p110α$^{-/-}$-eYFP lineage-tracing mice**

Female Pik3ca$^{fl/fl}$ mice on a C57BL/6 background were crossed with male Apoe$^{-/-}$ROSA26 STOP-floxeYFP$^{+/+}$Myh11-CreER$^{T22}$ on a C57BL/6 background (kindly provided by Gary K. Owens, University of Virginia, Charlottesville, Virginia, USA) to generate female Apoe$^{-/+}$Pik3ca$^{fl/wt}$ROSA*26* STOP-floxeYFP$^{+/-}$ and male Apoe$^{-/+}$Pik3ca$^{fl/wt}$ROSA*26* STOP-floxeYFP$^{+/-}$ Myh11-CreER$^{T2}$ mice, which were then inbred to produce male Apoe$^{+/+}$Pik3ca$^{fl/fl}$ROSA*26* STOP-floxeYFP$^{+/+}$Myh11-CreER$^{T2}$ and Apoe$^{+/+}$ Pik3ca$^{wt/wt}$ROSA*26* STOP-floxeYFP$^{+/+}$Myh11-CreER$^{T2}$ control mice.

Cre recombinase was activated in male mice with a series of 2-mg tamoxifen (Sigma, cat. no. T-5648) intraperitoneal injections in corn oil (Sigma, cat. no. C-8267; approximately 75 mg tamoxifen/kg body weight) once every 24 hours for a total of 5 consecutive days followed by a 9-day waiting period between the final injection and PPE induction to generate ism-p110α$^{+/+}$-eYFP and ism-p110α$^{-/-}$-eYFP mice.

**Cell processing for single-cell RNA sequencing.**

Six to eight ism-p110α$^{+/+}$-eYFP and ism-p110α$^{-/-}$-eYFP mice were euthanized by cervical dislocation 14 days after aneurysm induction. The mice were then perfused with 10 mL of PBS, and the infrarenal aorta was carefully stripped of fat and fascia before being removed from the animal. After removal from the animal, the tissue was placed in an enzyme cocktail containing 2 U/ml Liberase™ (Roche 05401119001), 0.1 mg/ml DNase I, and 60 U/ml hyaluronidase in RPMI-1640. After immersion in the digestion cocktail, the tissue was minced and placed in a 37 °C incubator for 1 hour. The digestion process was terminated by adding serum-containing medium; the cells were passed through a 70-µm strainer and centrifuged at 500 g for 5 minutes. The cells were resuspended in 200 µl of 1× DPBS with 0.1% FCS, and live YFP+ cells were sorted by flow cytometry after staining with the LIVE/DEAD Fixable Near IR (780) Viability Kit (Invitrogen L34992) using a BD FACSAria cell sorter. The gating strategy is shown in the **Supplementary Figure 5**.

**Single Cell Sequencing - Cell Isolation**

The sorted cell suspension was centrifuged at 350xg for 5 minutes at 4°C and resuspended in 1 ml of PBS after two washes with PBS. Cell counting and viability measurement were performed using acridine orange/propidium iodide double-staining on a Luna FX7 automated cell counter (Logos Biosystems). The cell suspension was loaded onto a SCOPE microfluidic chip from the GEXSCOPE™ Single Cell RNAseq Library Kit (Singleron Biotechnologies) to capture approximately 10,000 cells.

Transcriptome tagging was performed by adding paramagnetic beads conjugated with oligonucleotide barcodes and unique molecular identifiers (UMIs). Library construction was carried out according to the manufacturer's instructions.

The resulting single-cell RNA-seq libraries were sequenced using an Illumina NovaSeqX with a paired-end 150-bp configuration by GeneWiz Germany, GmbH (Leipzig, Germany). Sequencing reads were demultiplexed using Illumina's BaseCloud platform.

**Single Cell Sequencing - Transcriptome Preprocessing**

Raw sequencing reads were processed into gene expression matrices using CeleScope™ version 2.7.3 (www.github.com/singleron-RD/CeleScope; Singleron Biotechnologies).

FASTQ file barcode chemistry determination and barcode correction were performed before being demultiplexed and gene counts generated using the STARSolo single-cell pipeline with STAR version 2.7.11a [https://github.com/alexdobin/STAR] and STARsolo [https://github.com/cellgeni/STARsolo]. The mm10 draft of the mouse genome was used for mapping the reads and annotation was done with Ensembl version 99.

**Single Cell Sequencing - Data Processing and Analysis**

The transcriptional profiles of 12,912 and 15,860 cells for ism-p110⍰$^{+/+}$-eYFP and or ism-p110⍰$^{-/-}$-eYFP, respectively, were included in the analysis. Data was aligned using STARsolo (v2.7.11.a)[3] against the mm10 genome using default parameters. Ambient RNA was removed using cellbender (v0.3.2)[4] with a learning rate of 0.00005. The resulting data was processed using the scrapper R package (v1.6.3) using the sample as a blocking factor. Data wrangling was performed using the tidyverse (v1.3.0) and SingleCellExperiment (V1.34.0) R packages. Visualisations were generated using the tidyverse packages as well as the scatter (v1.40.1) R packages[5]. The contraction scoring for cells was calculated by taking the mean log-normalized expression of four core contractile genes (*Acta2*, *Cnn1*, *Myh11*, *Tagln*). For the proliferation scoring, the AUCell R package (v1.34) was used with mouse orthologs of cell-phase signatures[6,7,8]. Pathway analysis was performed with the decoupleR R package[9] using progeny pathways. [10]

**Cell Culture and treatment with siRNAs**

Murine SMCs were isolated from the thoracic and abdominal aorta by enzymatic dispersion as described.19 Animals were sacrificed via cervical dislocation. Subsequently, the aorta was dissected, removed, and incubated with 1 ml enzyme solution (1 mg/ml Collagenase Type I, 0.3 mg Elastase, 0.3 mg Trypsin inhibitor in DMEM / 20% FBS) for 15 minutes to detach the

adventitia. The remaining adventitia was removed using anatomic forceps, and the endothelium was gently removed by scraping the luminal surface. The aorta was cut into smaller pieces and incubated for further 90 minutes at 37°C in the enzyme solution to disintegrate the tissue and separate the SMCs. After centrifugation, the cell pellet was resuspended in DMEM culture medium containing 20% FCS and 1% Penicillin/Streptomycin. Cells were transferred into a well of a 24 well plate. After 2 days, most of the cells were attached to the plate. Culture medium was changed after three days. After reaching 80% confluence, cells were expanded. SMCs were grown in a 5% CO2 atmosphere at 37°C in DMEM supplemented with 100 U/ml penicillin, 100 µg/ml streptomycin, 1% nonessential amino acids (100X), and 10% FCS.
Murine aortic SMCs in 12-well plates were transfected with 150ng siRNA (FlexiTube GeneSolution GS56458 for Foxo1; Qiagen), respectively, according to the transfection-protocol of primary smooth muscle cells with siRNA using HiPerFect Transfection Reagent according to the manufacturer´s instructions (Qiagen).

**In vitro elastic fiber matrix analysis**
SMCs (1 x $10^5$) were seeded on 12 mm round coverslips in a 24-well plate and grown for seven days in 20% FCS/DMEM. Cells were fixed (4% PFA/PBS for 10 min at 37 °C) and blocked with 1% BSA/1xPBS. SMCs were permeabilized and primary antibodies diluted in blocking solution were added to the 24-well plate cover-slip and incubated for 1h at RT. After 3x washing in 1xPBS, incubation with the corresponding fluorescence-coupled secondary antibody (Anti-rabbit IgG Alexa Fluor 555 Conjugate #4413; 1:1000 diluted in 1% BSA/PBS + 0,3% TritonX100) was performed for 45 min at RT. In addition, DAPI (4,6-diamidine-2-phenylindole dihydrochloride) was added. The coverslips were then washed and mounted on slides with DAKO mount medium. Cells were washed with PBS containing 3% Triton X-100. Images were taken with a fluorescence (Axiophot, Zeiss) or confocal laser scanning microscope (SP5, Leica). Resultant SHG images were subjected to computational analyses using MATLAB software framework packages, CurveAlign and CT-FIRE as described.[11]

**Proliferation assay**
Real-time assessment of cell status was carried out with an IncuCyte Zoom System (Sartorius, Goettingen, Germany), as previously described.[12] Images were auto-collected and analyzed by using the IncuCyte software package. The cells were seeded into a 48-well plate (14000cells/well). All data are normalized with the cell confluence of 0h.

**BrdU incorporation**
DNA synthesis was measured by a BrdU-incorporation assay. Cells were cultured in 96-well plates (1 × $10^4$ cells per well) in DMEM containing 20% FCS for 24 hours, washed with PBS, and starved in DMEM for 24 hours. Subsequently, cells were preincubated with the specific FOXO1 inhibitor AS1842856 (1µM) or the selective p110α activator UCL-TRO-1938 (3-10µM) for 60 minutes followed by growth factor stimulation for 24 hours. BrdU incorporation was carried out according to the manufacturer’s specifications (Cell Proliferation ELISA, Roche)

with an incubation time of 5 hours as previously described.[13] Quantification was performed by measuring the absorbance using Power Wave 340 ELISA reader from Bio-Tek.

**Immunoblotting**

Quiescent SMCs were left resting or stimulated with growth factors, activator or inhibitor as indicated. Cells were washed twice with HEPES/sodium chloride (20 mM HEPES, pH 7.4, 150 mM NaCl), and then lysed in extraction buffer (10 mM Tris-HCl, pH 7.4, 5 mM EDTA, 50 mM NaCl, 50 mM NaF, 1% Triton X-100, 0.1% BSA, 20 µg/mL aprotinin, 2 mM Na3VO4, 1 mM PMSF). Lysates were centrifuged (20 minutes, 12,000g), and the supernatants were subjected to Western blot analysis. Similarly, tissue samples were homogenized in RIPA buffer. Protein concentration was assessed by either the Bradford Assay or NanoDrop quantification. Homogenates were suspended in 4 × SDS sample buffer. Equal amounts of protein lysates were run on SDS–PAGE and transferred to PVDF membranes. Blots were probed with the indicated antibodies.

**Immunohistochemistry & histochemistry**

Explanted aortae were fixed in 4% phosphate-buffered paraformaldehyde (Santa Cruz Biotechnology). The samples were embedded in paraffin and 3 µm sections were obtained from fixed tissue blocks. Morphometric analyses were performed using Image J analysis software (NIH). The mean intima/media was calculated in triplicates for each mouse by determining the area surrounded by the internal elastic lamina, the area surrounded by the external elastic lamina, and the lumen area. Nuclear staining was performed with Harries hematoxylin following standardprocedures. Immunohistochemistry was performed using antibodies directly against the phosphorylated AKT (Cell Signaling Technology, USA, catalog 3777) with 3,3'-diaminobenzidine as a substrate (DAB Substrate Kit, Linaris) as well as calponin (Cell Signaling Technology, USA, catalog 17819) and $\alpha$-sm-actin (Cell Signaling Technology, USA, catalog 19245) with Vector®-VIP kit, Vector Laboratories, USA. Negative controls were performed with the omission of the primary antibody. Hematoxylin was used to counterstain. Tissue sections were stained for PCNA (Santa Cruz Biotechnology, sc-7907) to detect proliferation.

Aortic sections were stained with Picro-sirius red (Picro Sirius Red Stain Kit (Connective Tissue Stain), Abcam) for 1 h at room temperature to visualize collagen fibers and washed with 1% acetic acid twice for 10 min at room temperature. The picro sirius red staining was viewed using standard light microscopy. Measurements were made using the QuPath image analysis software to measure the amount of red (collagen) and yellow (muscle fibers, cytoplasm) color relative to the aortic area.

For elastin, aortic sections were stained with Verhoeff's hematoxylin for 1 h, differentiated in 2% ferric chloride for 2 min and counterstained with Van Gieson solution for 5 min (all at room temperature) to identify elastic fibers in the aortic tissue. For analysis, a scoring system between 1 and 4 was used as described elsewhere[14]: (1) intact elastin layer; (2) mild degradation of elastin and some laminar disruption or rupture; (3) moderate elastin degradation and multiple interruptions or breaks in the lamina; (4) severe elastin breakdown or loss or aortic rupture.

**Second harmonic generation**

For collagen detection, 10 µm cross-sections of paraffin-embedded aortas were dewaxed, and second harmonic generation microscopy (SHG) of aortic collagens was obtained by using a Leica TCS SP8 microscopy (Leica microsystems, Wetzlar Germany) with a Chameleon Vision IIlaser. SHG and TPEF were excited at 810 nm and imaged with an IR Apo L25x/0.95 W objective lens.

**Real-time qPCR**

Briefly, aortae were isolated and homogenized using Precellys Lysing Kit (Bertin technologies) and Qiashredder (Qiagen). mRNA from aortae was isolated with RNeasy micro kit (QIAGEN). cDNA was prepared with SuperScript III first-strand synthesis system (Invitrogen). qRT–PCR of *Acta2, Cnn1, Klf4, Eln, Col1a1, Mmp2, Mmp9, Cd68, Foxo1,* and *Gapdh* genes was performed using TaqMan gene expression assays (Applied Biosystems).

**Materials and antibodies**

Chemicals were obtained from Sigma-Aldrich Chemie GmbH. Rat tail collagen type I was obtained from BD Biosciences, elastase from Serva Electrophoresis GmbH, collagenase from Sigma-Aldrich Chemie GmbH, UCL-TRO-1938 from MedChemtronica, and AS1842856 from Sigma-Aldrich (Merck). The antibody against RasGAP (clone 69.3) was a gift from Andrius Kazlauskas (University of Illinois, Chicago, U.S.) and the fibrillin-1 antibody was kindly provided by G. Sengle (University Hospital of Cologne, Germany). The phospho-Akt Thr308 (catalog 13038), phospho-AKT1 (catalog 9018), phospho-AKT2 (catalog 8599), phospho-GSK3β (catalog 9322), phospho-FOXO 1&3 (catalog 9496), phospho-FOXO4 (catalog 9471), p110α (catalog 4255), calponin (catalog 17819), PCNA (catalog 13110), and anti-rabbit IgG (H+L), F(ab')2 Fragment (Alexa Fluor® 555 Conjugate) (catalog 4413) antibodies were obtained from Cell Signaling Technology; α-sm-actin (catalog 53141) antibody from Santa Cruz Biotechnology; MOMA-2 (ab3345), 1β-actin (catalog ab2827) and tropoelastin (catalog ab21600) from Abcam; and Myosin-heavy-chain (catalog MAB4470) from R&D systems. PDGF-BB (catalog C-63022), Insulin (catalog C-60840) were purchased from Promo Cell, and AS1842856 from Sigma-Aldrich.

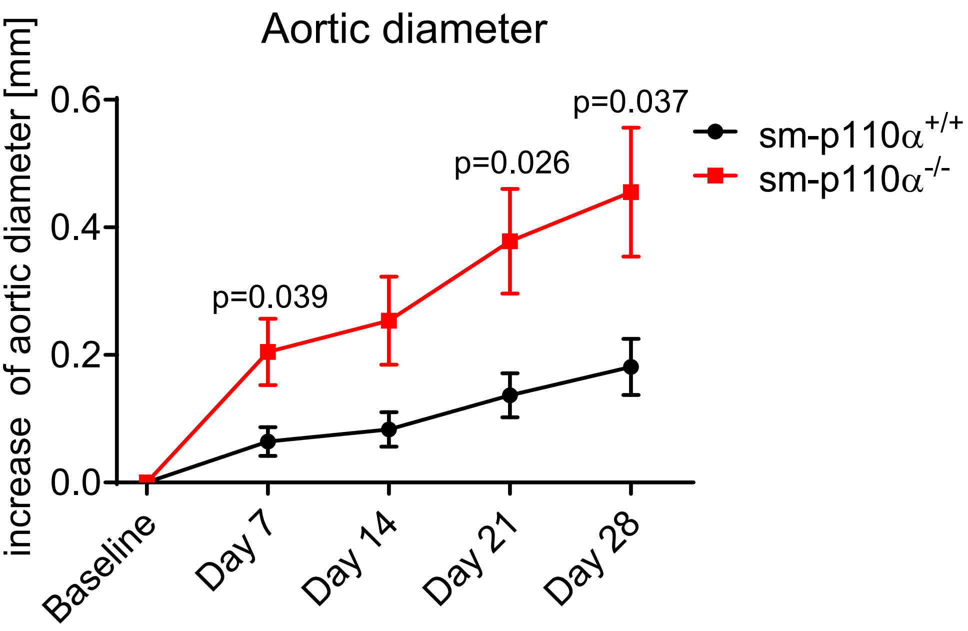


**Supplementary Figure 1.** Increase of the aortic diameter 0, 7, 14, 21 and 28 days after PPE intervention in mm. n=9-10. Data is presented as means ± sem. P value was calculated mixed-effects analysis followed by Bonferroni correction.

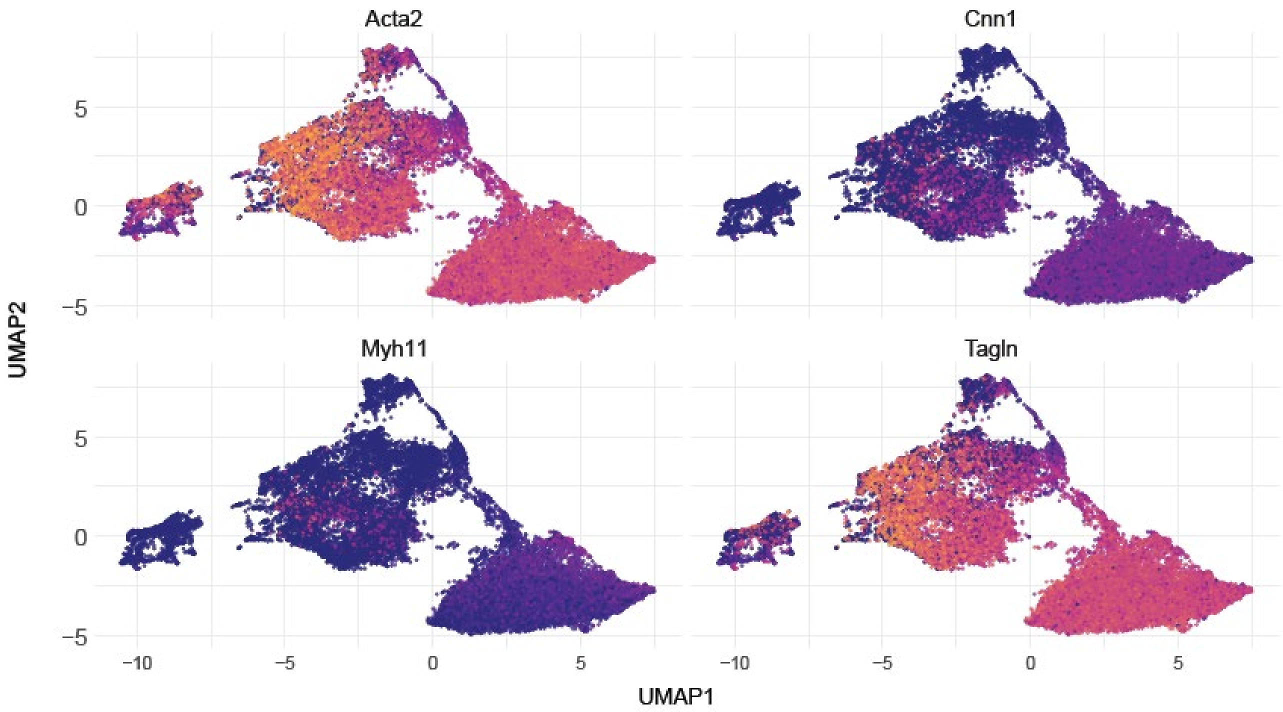


**Supplementary Figure 2.** Feature plots demonstrate the expression of contractile genes (*Acta2, Cnn1, Tagln, Myh11)* in 4 SMC type clusters. Color scale indicates log2 expression values, from 0 (low) to 8 (high).

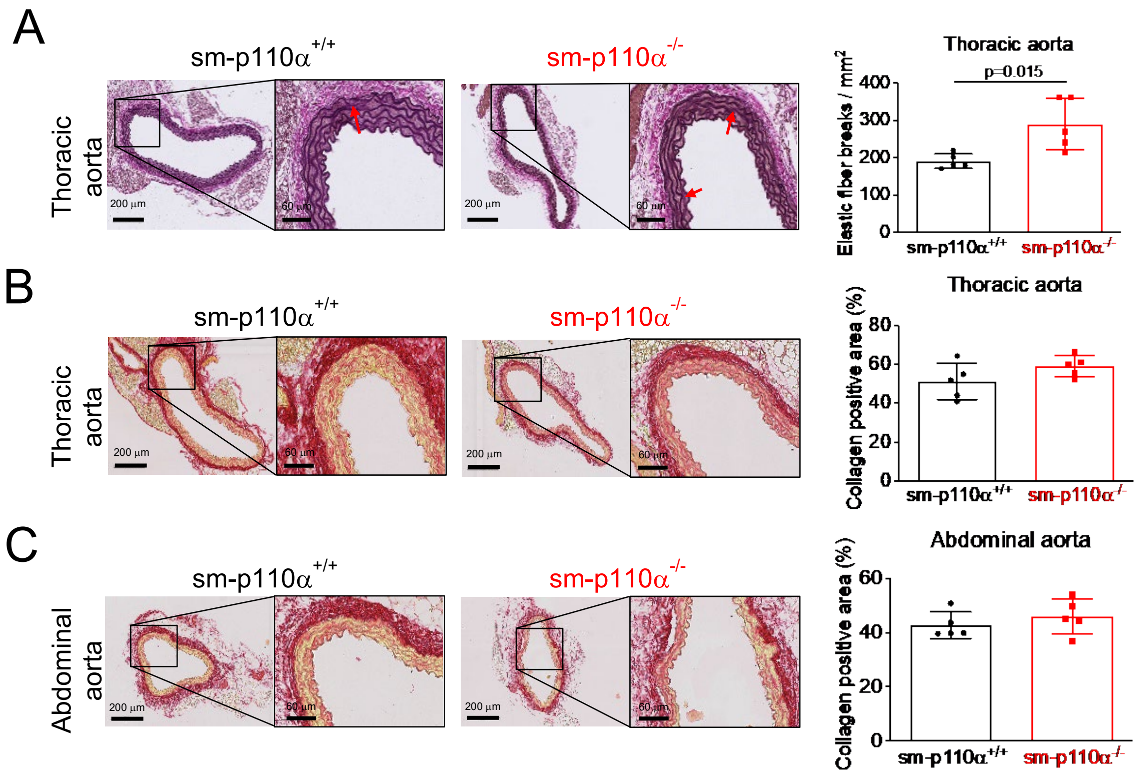


**Supplementary Figure 3. A** Representative images of elastic van Gieson staining (100 x and 400 x magnification) in cross section from thoracic aorta from 13 weeks old sm-p110α$^{+/+}$ versus sm-p110α$^{-/-}$ mice under basal conditions and quantifications of elastic fiber breaks, respectively. Scale bars 200 µm (overview) and 60 µm (zoom). The number of elastic fiber breaks was counted per cross section (n=5, respectively) and shown per mm$^2$ media area. **B, C** Representative Picro Sirius Red staining for fibrotic collagen of cross-sections from (**B**) thoracic and (**C**) abdominal aortae from 13 weeks old sm-p110α$^{+/+}$ and sm-p110α$^{-/-}$ mice as indicated. Scale bars 200 µm (overview) and 60 µm (zoom). Collagen was quantified (n=3-5). *P* value was calculated using an unpaired two-tailed Student's t-test. Data is presented as means ± S.D.

A

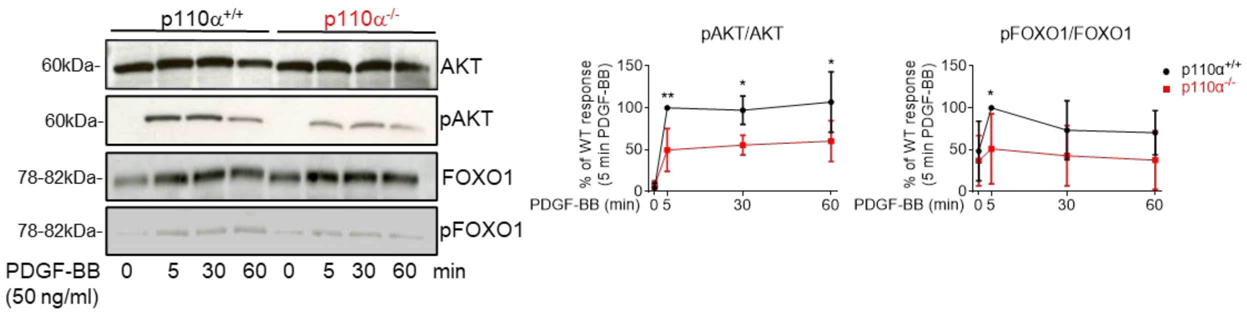


B

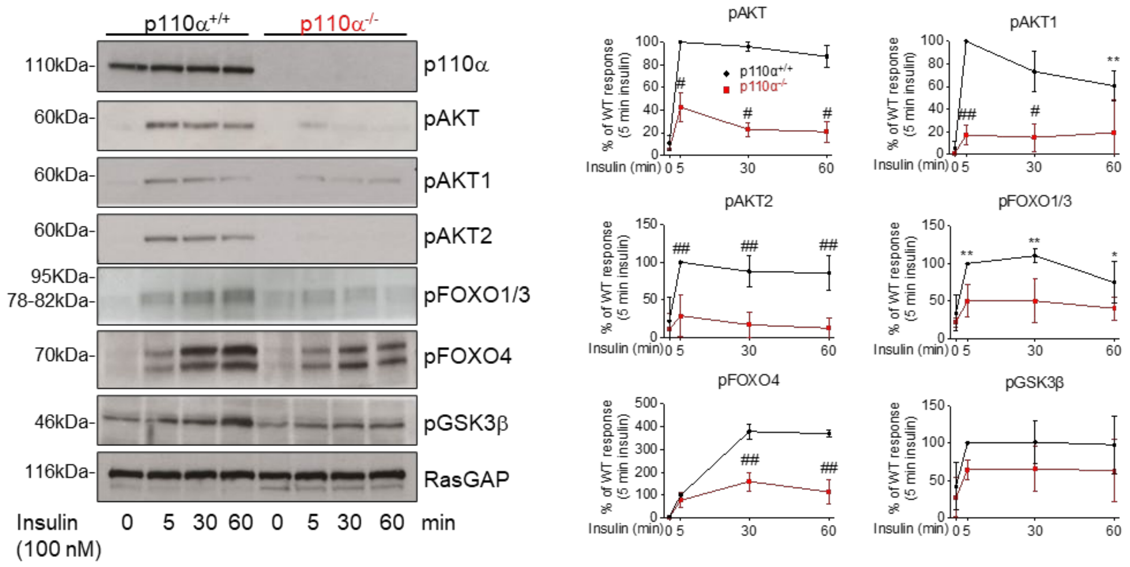


**Supplementary Figure 4: p110α signaling in insulin stimulated SMCs. A, B** Representative Western blots of cultured PDGF-BB (50 ng/ml) (**A**) or insulin (100 nM) (**B**) stimulated aortic SMCs detecting phosphorylated signaling proteins downstream of p110α as well as loading controls as indicated. Densitometrical analyses are shown on the right. (n=3-4). *P* values were calculated using a 2-way ANOVA followed by Bonferroni correction. *$p < 0.05$, **$p < 0.01$, $^{\#}p<0.001$, $^{\#\#}p<0.0001$. Data is presented as mean ± S.D.

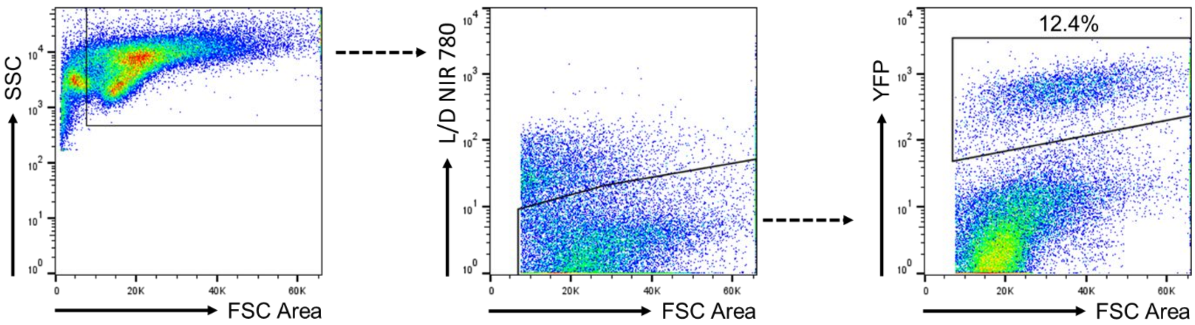


**Supplementary Figure 5**. Gating strategy to isolate YFP+ cells of SMC origin. Flow cytometry of single cell suspension from infrarenal aortas of ism-p110$\alpha^{-/-}$-eYFP mice. Gating strategy includes: Forward vs Side Scatter to exclude debris and leukocytes, Side Scatter vs LIVE/DEAD Fixable Near IR (780) Viability Kit (L/D NIR 780) to identify live cells, and Side Scatter vs YFP to isolate YFP+ cells of SMC origin (representative experiment, n = 6).

| Category | Details | Counts / Values |
|---|---|---|
| Age (mean), years | | 65 |
| Gender | Male | 12 |
| | Female | 4 |
| Hypertension | Yes | 15 |
| | No | 1 |
| Diabetes | Yes | 2 |
| | No | 14 |
| Renal insufficiency | Yes | 9 |
| | No | 4 |
| | Not Available (n/a) | 3 |
| Statin | Yes | 11 |
| | No | 5 |

**Supplementary Table 1.** Patient Information
